# A chromosome-level genome assembly for the Pacific oyster (*Crassostrea gigas*)

**DOI:** 10.1101/2020.09.25.313494

**Authors:** Carolina Peñaloza, Alejandro P. Gutierrez, Lel Eory, Shan Wang, Ximing Guo, Alan L. Archibald, Tim P. Bean, Ross D. Houston

## Abstract

The Pacific oyster (*Crassostrea gigas*) is a marine bivalve species with vital roles in coastal ecosystems and aquaculture globally. While extensive genomic tools are available for *C. gigas*, highly contiguous reference genomes are required to support both fundamental and applied research. In the current study, high coverage long and short read sequence data generated on Pacific Biosciences and Illumina platforms from a single female individual specimen was used to generate an initial assembly, which was then scaffolded into 10 pseudo chromosomes using both Hi-C sequencing and a high density SNP linkage map. The final assembly has a scaffold N50 of 58.4 Mb and a contig N50 of 1.8 Mb, representing a step advance on the previously published *C. gigas* assembly. The new assembly was annotated using Pacific Biosciences Iso-Seq and Illumina RNA-Seq data, identifying 30K putative protein coding genes, with an average of 3.9 transcripts per gene. Annotation of putative repeat elements highlighted an inverse relationship with gene density, and identified putative centromeres of the metacentric chromosomes. An enrichment of *Helitron* rolling circle transponsable elements was observed, suggesting their potential role in shaping the evolution of the *C. gigas* genome. This new chromosome-level assembly will be an enabling resource for genetics and genomics studies to support fundamental insight into bivalve biology, as well as for genetic improvement of *C. gigas* in aquaculture breeding programmes.

## Background

The Pacific oyster, *Crassostrea gigas* (Thunberg, 1793) (NCBI:txid29159), also referred to as *Magallana gigas* by some authors [1, 2], is a keystone ecosystem and aquaculture species [3]. Although native to the Pacific coast of Northeast Asia [4], *C. gigas* has been introduced to all continents, except Antarctica, for farming purposes [5-9]. The intensive human-mediated spread of Pacific oysters was mainly catalysed by the collapse of the fishery and culture of native oyster stocks due to disease, and the need to supplement depleted stocks [10, 11]. Most of these initiatives had far-reaching effects on the global distribution of Pacific oysters, since several self-sustaining populations became established in the wild [12, 13]. As a result, *C. gigas* is now one of the most highly produced aquaculture species globally, and a conspicuous invasive species in many countries [14].

The extent of genetic and genomic resources developed for Pacific oysters are unparalleled among molluscs and other lower invertebrates [15]. Hence, they are often used as model organisms to represent Lophotrochozoa, a major clade showing the greatest diversity of body plans among Bilaterians [16-18]. These resources have also been applied to enhance aquaculture production, with early technological advances in *C. gigas* focused on developing techniques to improve production through ploidy manipulation [19, 20], which later allowed the creation of the first tetraploid and triploid oyster stocks [21]. Advances in DNA sequencing technologies led to rapid additional resource development for this species, including extensive transcriptome datasets [22-26], linkage maps using microsatellite and more recently SNP markers [27, 28], and medium and high density SNP arrays [29, 30]. These tools have become valuable genomic resources to enhance genetic improvement of production traits, such as growth and disease resistance [31, 32]. Nevertheless, a key resource for enabling genetics and genomic research in a given species is a high quality reference genome. Zhang, Fang [33] published the first draft reference genome assembly for *C. gigas* using a whole genome shotgun sequencing approach and short read Illumina sequenced data. Interrogation of the reference genome data pointed to gene expansion as a likely factor explaining the adaptation of *C. gigas* to challenging marine environments, a finding that has been mirrored in a number of subsequent reference genome studies for bivalve shellfish (reviewed in [34]). Although a major achievement, and indeed one of the first genome assemblies for a molluscan species, the publicly available reference genome is highly fragmented (GenBank accession number GCA_000297895.2, 26,965 contigs, contig N50 = 42.3 Kb). Moreover, the previous version of this assembly (GCA_000297895.1) contains many misplaced and chimeric scaffolds as revealed by alignment with linkage maps [28, 30]. These issues are likely to derive from a combination of both biological factors, such as the high levels of genome heterozygosity, and technical factors, such as the reliance on short-read sequencing available at the time [33]. Therefore, highly contiguous and accurate reference genome assemblies would represent valuable resources for enabling genetics and genomic research in this keystone species.

In the current study, an improved (chromosome-level) assembly was developed for *C. gigas* by harnessing high coverage Pacific Biosciences (PacBio) long-read sequencing (80X), alongside accurate Illumina short read data (50x). The assembly was then scaffolded to chromosome level using both Hi-C sequencing and a high-density SNP linkage map, and the genome was annotated based on both Illumina and PacBio transcript sequencing. This improved reference genome assembly represents a step forward towards improving our understanding of fundamental biological and evolutionary questions, and the genetic improvement of important aquaculture production traits via genomics-enabled breeding.

### Sample collection and sequencing

A single female individual collected in 2017 from Guernsey Sea Farms (Guernsey, UK) was used for whole-genome resequencing with the PacBio Sequel (Pacific Biosciences, Menlo Park, CA, USA) and the HiSeq X (Illumina Inc.; San Diego, CA, USA) platforms. High quality dsDNA was isolated from ethanol-preserved gill tissue using a cetyl trimethylammonium bromide (CTAB) method. The quality of the DNA extraction was verified by the NanoDrop A260/280 and 260 /230 ratios (ND-1000) and a fluorescence-based electrophoresis on a 2200 TapeStation System (Agilent Technologies, USA). Using this purified DNA, three different types of libraries were prepared to generate the sequencing data used for the assembly of the *C. gigas* genome. The first set of libraries were generated to obtain long PacBio reads and develop an initial *de novo* assembly. Two SMRTbell® libraries (chemistry v3.0) were prepared and sequenced by Edinburgh Genomics (University of Edinburgh, UK) across 13 SMRT cells of a PacBio Sequel system. A total of ∼55 Gb of raw bases with an N50 length of 12,777 bp were produced (Supplementary Figure S1). Second, a paired-end sequencing library of 300 bp insert size was prepared from the same individual and then used for (i) sequence error correction and (ii) quality assessment of the draft genome assembly. This library was produced by Edinburgh Genomics using the TruSeq DNA Nano gel free library kit (Illumina) and then sequenced on a HiSeq X platform (2 x 150 bp paired-end reads). Approximately 210 million short reads were obtained after quality filtering (average BQ>15 over 5 bp) and adapter removal with Trimmomatic v0.38 [35]. Thirdly, a Hi-C library was generated with the purpose of scaffolding the assembly into large pseudo-chromosomes. Libraries were prepared using the Dovetail^™^ Hi-C Library Preparation Kit, following the manufacturer’s protocol (Dovetail™ Hi-C Kit Manual v.1.03). This final library was sequenced on an Illumina HiSeq X platform (2 x 150 bp), and resulted in 500 million read pairs.

Total RNA was extracted from two additional individual oysters (also from Guernsey Sea Farms, Guernsey, UK), a male and a female, from six distinct tissues (gill, mantle, stomach, heart, adductor muscle and gonads (ovaries and testis)). Full-length transcripts were isolated from the tissue samples using a combination of the TRIzol (Invitrogen) and the RNeasy plus minikit (Qiagen) protocols, with the inclusion of a DNAse treatment step. RNA quality was assessed using the Nanodrop ND-1000 and the Agilent 2200 TapeStation instruments. RNA extracts were quantified using a Qubit^™^ RNA assay kit (Thermo Fisher, Waltham, MA, US), and then combined in equimolar quantities into a single pool for sequencing. The final RNA-pool was used to obtain full-length cDNA sequences using the TeloPrime Full-Length cDNA Amplification Kit V2 (Lexogen). cDNA was then sequenced across three SMRT cells of a PacBio Sequel platform at the Dresden-concept Genome Center DcGC (Germany). A total of 178 Gb of data comprising 1.6 million transcripts with a mean length of 1.3 kb were generated for gene annotation.

### Genome features

Due to the differences in genome size estimates reported in the literature for *C. gigas* [15, 33], the DNA content of the Pacific oyster genome was also estimated in the current study. To this end, the average genome size was estimated for the sequenced female using the k-mer method [36] and flow cytometry [37]. For the k-mer based approach, quality-filtered Illumina reads (150 bp length) were used to count the frequency of different *k*-mer sizes, ranging from 15 to 23, using Jellyfish v2.1.3 [36]. All *k* values evaluated showed a clear bimodal distribution, with peaks occurring at a read depth of 19X and 37X (Supplementary Figure S2). The k-mer frequency plots obtained are characteristic of species with highly heterozygous genomes [38]. From the k-mer based analysis (k-mer = 21), the *C. gigas* genome size was estimated at 534 Mb. For the genome size estimation by flow cytometry, Pacific oyster nuclei were isolated and stained with propidium iodide. Two species were used as internal standards for the assay, fruit fly (*Drosophila melanogaster*) and zebra fish (*Danio rerio*). According to flow cytometry measurements, the genome size of the female oyster sequenced in the current study was estimated at 640 Mb. Due to the different genome size estimates obtained by the two methods, the midpoint – i.e. ∼590 Mb -was used to calculate the predicted sequencing yield and anticipated length for *de novo* genome assembly. The heterozygosity of the Pacific oyster genome was assessed with GenomeScope v2.0 [39], based on the quality filtered Illumina reads. A heterozygosity rate of 3% was estimated from the 21-mer based assessment of the oyster genome (Supplementary Figure S3). This value is higher than the 1.3% previously reported for this species [33], but is likely explained by the fact that the authors used an inbred individual for genome assembly, whereas in this study an outbred female was sequenced. Although high, the heterozygosity value is in the range with those reported for other bivalve molluscs (e.g. 2.4% in the quagga mussel [40]).

### Genome assembly

The PacBio reads were first assembled into contigs using Canu v1.8 [41] at near default parameters (corrected error rate = 0.045 and raw error rate = 0.300). Contigs were polished with one round of Arrow [42] followed by an additional round of polishing with Pilon [43], after alignment of the post-quality filtered Illumina reads with Minimap2 [44]. Compared with the genome size estimate of 590 Mb, the initially assembled version of the genome was approximately two times larger than expected, yielding 6,368 contigs, a total length of ∼1.2 Gb, and an N50 length of 0.46 Mb. These results can be explained by the high frequency of highly divergent haplotypes in the *C. gigas* genome, a feature that has also been observed in the process of creating genome assemblies for other molluscan species [45, 46]. Whilst the size of the assembled sequence could indicate that the high level of heterozygosity had allowed the resolution of the two haplotypes present, we sought to establish a high quality pseudo-haploid genome as a reference. To assess the level of duplication in the initial assembly, a BUSCO analysis was performed [47]. By searching against the metazoa_odb9 database using sea hare as a reference species, 791 BUSCO genes (80.9%) were found to be duplicated. To remove potentially redundant contigs by retaining only one variant of a pair of divergent haplotypes, two independent approaches were taken. First, the short read data were used to identify and reassign putative haplotigs with the Purge Haplotigs pipeline (-l 5, -m 38, -h 90) [48]. Secondly, an all-versus-all contig mapping was performed on the repeat masked assembly with minimap2 v.2.2.15 [44]. Contigs were ordered based on their length and matching contigs which mapped at least 30% of their length and longer than 10kbp were removed as potential haplotigs. The reference sequence and the mapping sequences were all removed before the next iteration. The lists of curated contigs obtained independently from both methods were compared and the common contigs then selected for an additional round of haplotig purging. This approach resulted in a significant reduction in the number of contigs to 1,235, which were retained for scaffolding.

### Chromosome-level assembly using Hi-C and linkage map data

To generate a chromosome level assembly for *C. gigas*, Hi-C proximity ligation [49] data were used to order and orientate the contigs along chromosomes. The scaffolding process was carried out by Dovetail Genomics (Santa Cruz, CA, USA) using the Dovetail^™^ Hi-C library reads to connect and order the input set of contigs. After scaffolding with HiRise v2.1.7 [50], the assembled genome sequence initially comprised a total of ∼ 633 Mb, with a scaffold and contig N50 of 57 and 0.7 Mb and, respectively. A high fraction of the assembled sequences (>92%) was contained in only 11 super-scaffolds (Figure 1). However, Pacific oysters have 10 pairs of chromosomes [51]. A high-density linkage map [27] was used to anchor the super-scaffolds into chromosomes. SNP probes were mapped to the reference genome assembly using BWA v0.78 [52]. Of the 20,353 markers on the genetic map, 17,747 mapped to a chromosome-level scaffold with a MAPQ above 16. The integration of genetic-linkage information enabled the anchoring of two super-scaffolds onto a single linkage group (LG2), resulting in an assembly with 10 major scaffolds that represent all oyster chromosomes (Figure 2). Gaps were closed with PBJelly [53] and again error corrected using the short read Illumina data using Pilon [43]. From the remaining set of unplaced scaffolds, regions of low sequence accuracy were identified based on short read coverage, following [54]. Briefly, the median read-depth per 1,000 bp (non-overlapping) windows was calculated, after GC-content normalization. Scaffolds with >70% of windows showing a median coverage 2SD above or below the mean were removed from the analysis. All unplaced contigs and scaffolds showing significant sequence identity with the Iso-Seq data were added to the primary set.

**Figure 1.**
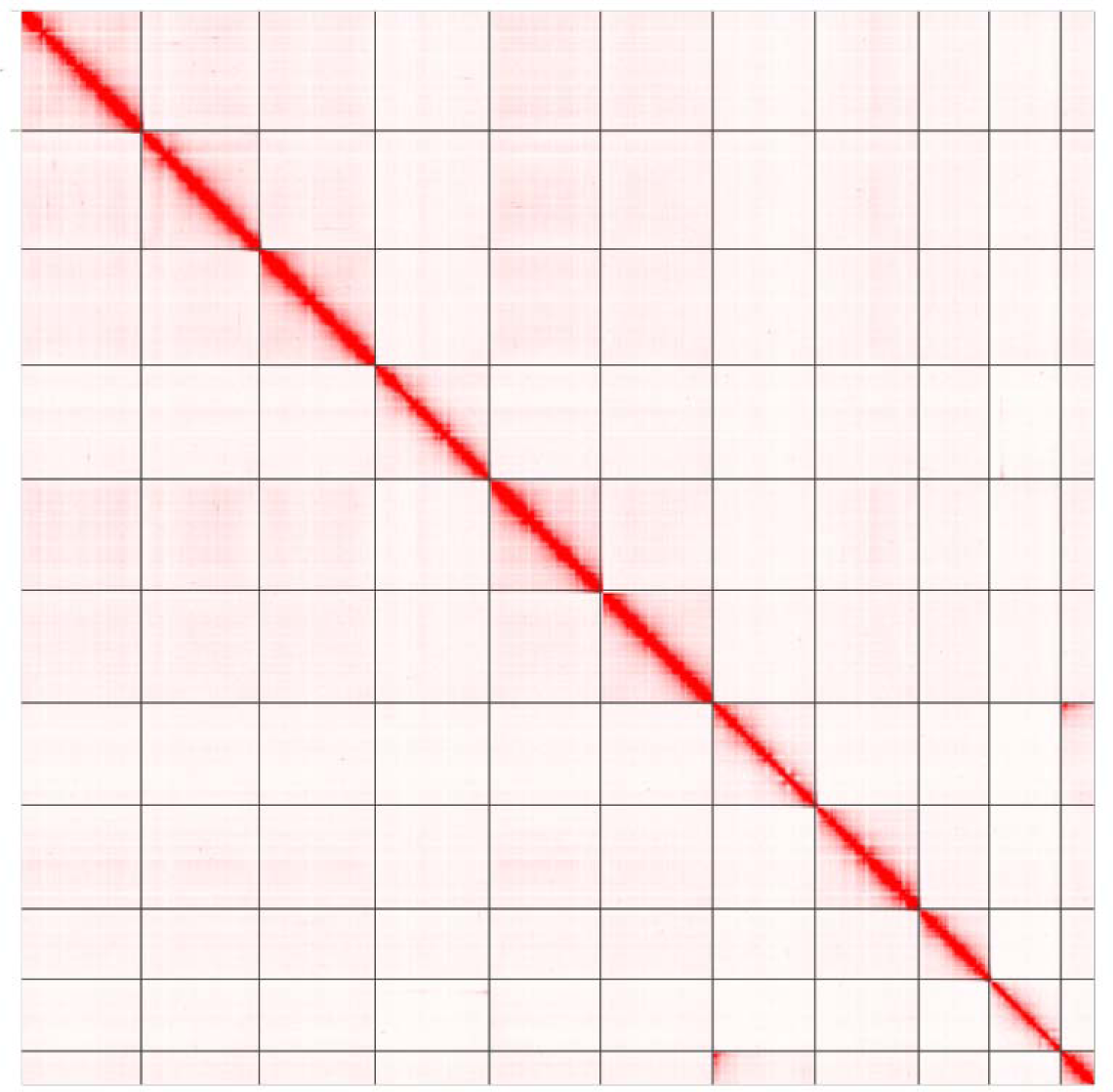
Hi-C interaction analysis depicting the 11 super-scaffolds obtained after using the HiRise™ scaffolding software. Contact map is visualized using Juicebox v1.11.08 [55].

**Figure 2.**
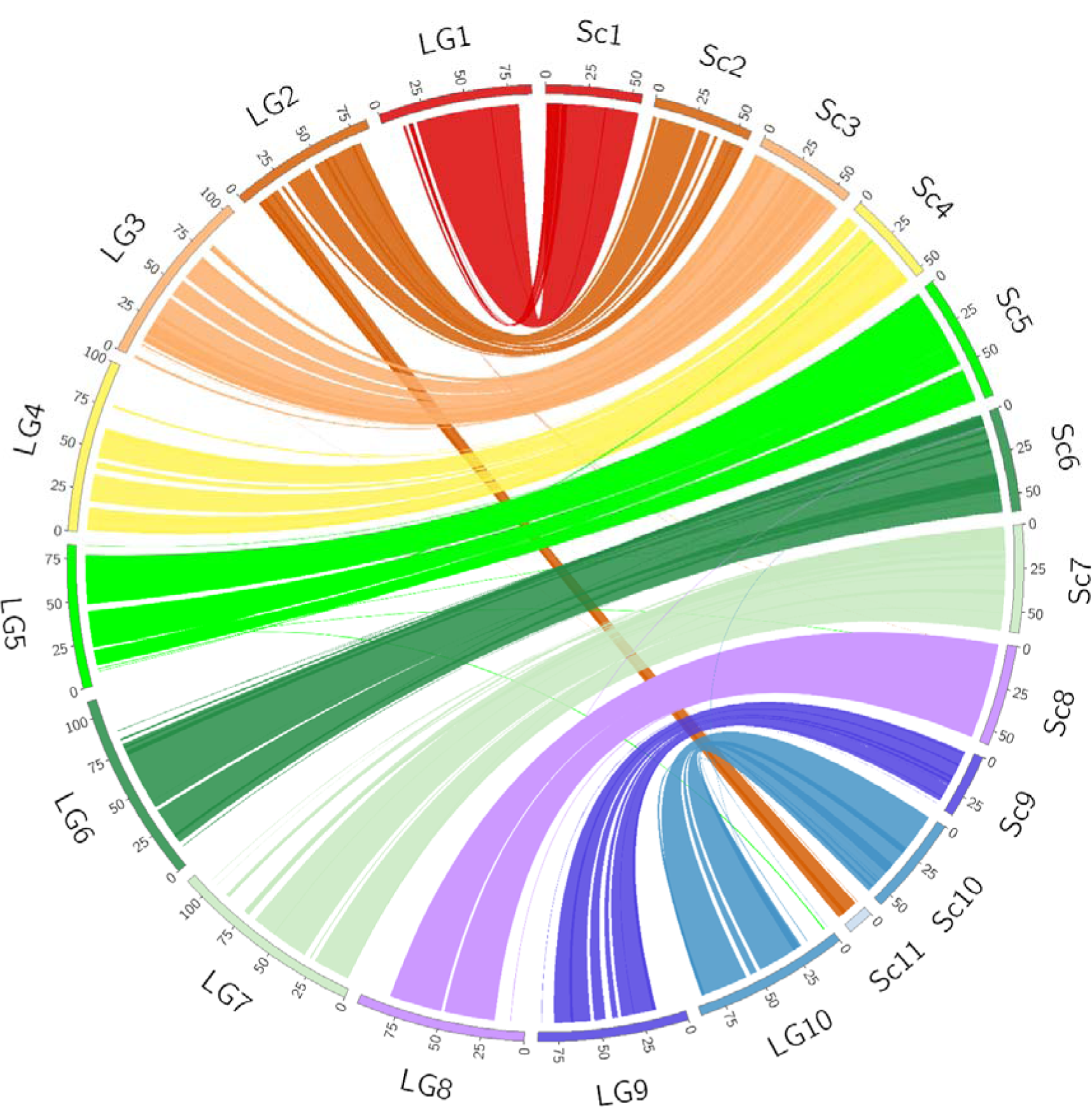
A high concordance between the chromosome-level scaffolds and a high-density linkage map allowed the anchoring of two scaffolds (Sc2 and Sc11) to a single linkage group 2 (LG2). Ticks in each linkage group or scaffold indicate lengths in 25 Mb. Scaffold (Sc) unit lengths are in Mb. Linkage group (LG) units of distance are expressed in cM. Plot generated using Circos v0.69-8 [56].

The final Pacific oyster assembly (GenBank accession number GCA_902806645.1) contains the ten expected chromosomes and 226 unplaced scaffolds, with a total N50 of 58.4 Mb and 1.8 Mb for scaffold and contig lengths, respectively (Table 1). This final assembly is 647 Mb in size, with the chromosome-level scaffolds represented in 589 Mb of sequence. This represents a step improvement over the previous version of the *C. gigas* reference genome [33], and other oyster assemblies [46]. However, it should be noted that a separate chromosome-level reference genome assembly is available in GenBank (accession number GCA_011032805.1) from the Institute of Oceanology, Chinese Academy of Sciences. This assembly is slightly shorter at 587 Mb, has a similar scaffold N50 of 61.0 Mb, and a higher contig N50 of 3.1 Mb. Future comparisons between these two high quality assemblies will be important to evaluate their consistency and ensure uniform use of nomenclature to describe chromosomes. Furthermore, it is expected that additional high quality reference genome assemblies will become available for this species, and the availability of multiple assemblies is advantageous for *C. gigas* as a species with high levels of intra- and inter-population genetic diversity [15]. To aid with the coordination of this assembly with existing and future assemblies, the ten large scaffolds in the current assembly were aligned with the karyotype using FISH probes corresponding to BAC clones (Supplementary Note A). The correspondence between the nomenclature of the linkage groups and scaffolds assembled in the current study, and the chromosome number of the karyotype is given in Supplementary Table 1. This information should enable consistency in nomenclature when describing multiple genome assemblies for this species in the future.

**Table 1.**
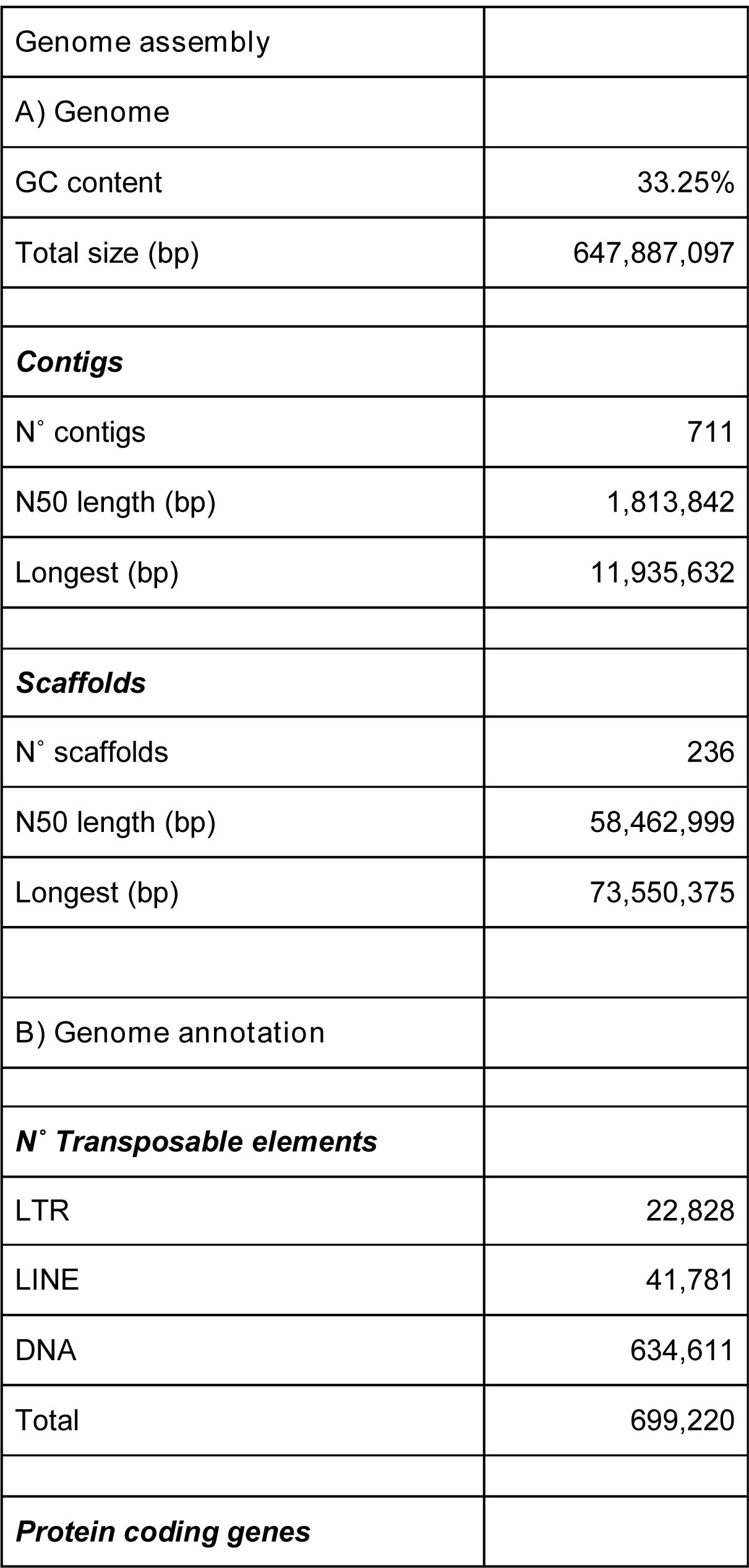
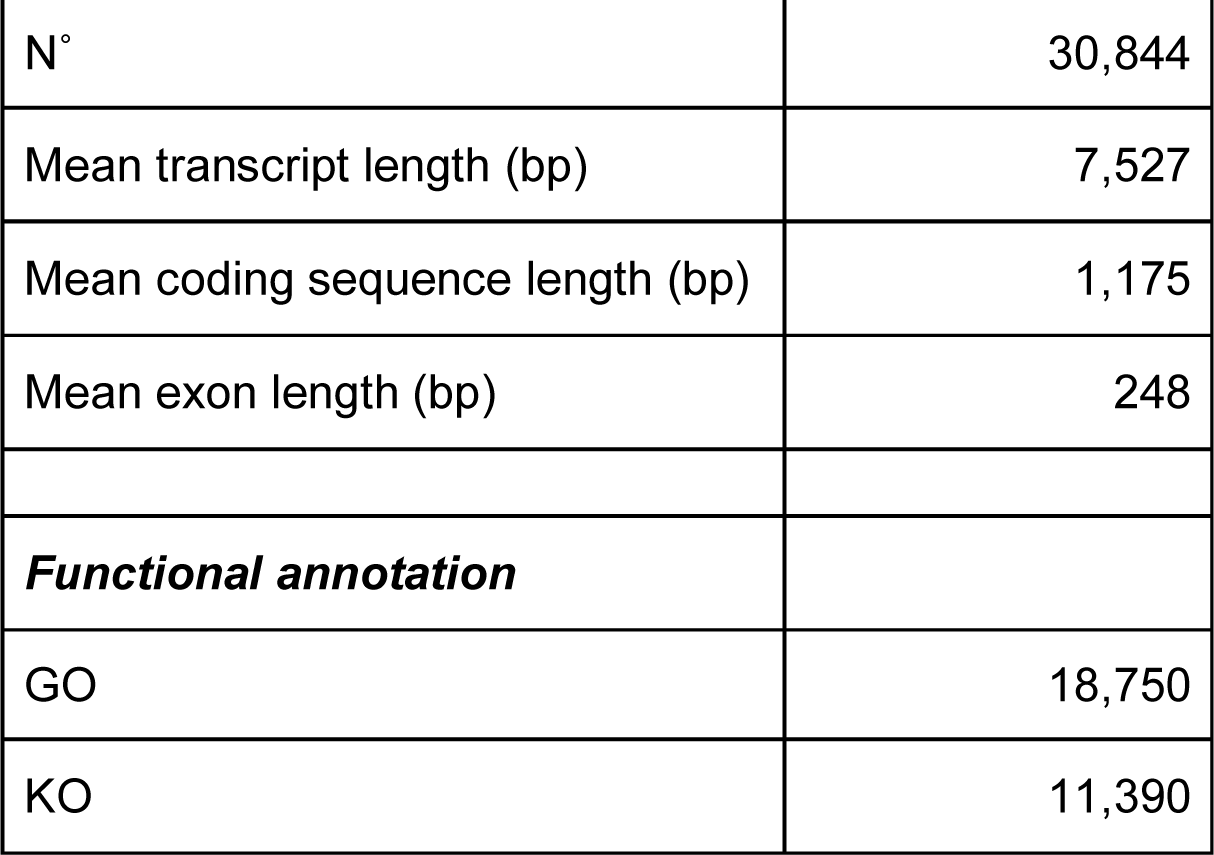
Genome assembly statistics and annotation of *C. gigas*.

### Quality assessment of reference genome

Firstly, the assembled *C. gigas* genome was screened for contaminant DNA with BlobTools v1 [57]. All scaffolds and contigs had a top hit match to *C. gigas* (Supplementary Figure S5). Second, to assess the completeness of the assembled genome, a BUSCO analysis was performed. From the curated list of single copy genes, 934 (95.5%) were found in the assembly, of which 917 (93.8%) were single-copy genes, 17 (1.7%) were duplicated and 38 (3.9%) were missing. Finally, to evaluate the accuracy of the reconstructed *C. gigas* genome, structural variants were called with Sniffles [58], after alignment of the PacBio raw reads with ngmlr v0.2.7. Variants with a minimum size of 50 bp for which the ratio of high quality reads for the assembly (reference) variant was below 0.2 were considered assembly errors (Table S2).

### Genome annotation

Genome annotation was carried out using long-read PacBio Iso-Seq data from six tissues and the Illumina short-read RNA-Seq data from Zhang [33]. Short-read data was mapped to the reference assembly with STAR v.2.5.1b [59]. Transcript models were created by BRAKER v.2.1.5 [60] using only the paired-end RNA-seq datasets (see Supplementary Table S3). Multi exon transcripts that were expressed in at least two tissues at an expression level over 1TPM were retained. Iso-Seq raw sub-reads were processed with SMRT Link v7.0 (Pacific Biosciences) to obtain Circular Consensus Sequences (CCS) using a ‘--min-rq of 0.9’. The Iso-Seq CCS reads were mapped with minimap2 v.2.16 [44] and the transcript models were called using the TAMA package [61] (see Supplementary Note B). Protein-coding transcripts and translation start and end positions were predicted by mapping known protein sequences from UniRef90 [62] to the oyster transcripts by Diamond v.0.9.31 [63]. Those models that contained a frameshift within the coding sequence were classified as pseudogenes.

The final annotation of the assembled *C. gigas* genome contains 35,527 genes, of which 30,844 are protein coding, 4,001 represent non-coding RNA genes and 682 were classified as pseudogenes. Among the protein coding genes, 14,293 (49%) contained putative alternative spliced transcripts, with an average of 3.9 transcripts per gene. The gene models predicted for *C. gigas* were functionally annotated using the Blast2GO pipeline [64], and KEGG orthology (KO) groups were assigned using KOBAS v2.0 [65]. Approximately, 18,750 (61%) of the predicted protein coding genes were assigned functional labels (Table 1). This reference genome assembly has also been annotated by the NCBI annotation team, who used the extensive short read transcriptome data available for *C. gigas* to annotate 38,296 genes (31,371 protein coding, 6,837 non-coding, 88 pseudogenes) and a total of 73,946 transcripts [66].

### Repeat element annotation

Known Pacific oyster specific repeat sequences were identified in the genome assembly using RepeatMasker v.4.0.7 [67] with a combined repeat database (Dfam_Consensus-20170127 and RepBase-20170127) [68, 69] with parameters ‘-s -species “Crassostrea gigas” -e ncbi’. Besides the 972 repeat families contained in the RepeatMasker library an additional 1,827 novel repeat families were identified by RepeatModeler v.1.0.11 [70]. This novel repeat library was used to identify the location of novel elements in the newly built assembly. For comparison, an exact same search was performed on the older version of the *C. gigas* genome assembly (GenBank assembly accession GCA_000297895.2).

Overall, a higher number of repetitive elements were identified in our assembly compared to the previous genome assembly (Figure S6). Nearly 43% of the Pacific oyster genome was represented by repeat elements. Repetitive sequences were distributed unevenly along the *C. gigas* chromosomes. In general, an inverse relationship between the total number of repeat elements and gene density was observed (Figure 3 d-e). Among the different classes of repeat elements, significant negative correlations were found between gene density and (i) retrotransposons of the LTR type (corr = −0.61; P = 2.2 x 10^−16^), (ii) Non-LTR retrotransposons (corr = −0.28; P = 5.4 × 10^−7^), (iii) satellite DNA (corr = −0.29; P = 4.5 × 10^−7^), (iv) simple repeats (corr = −0.33; P = 4.7 × 10^−9^), and (v) DNA transposons (corr = −0.59; P = 2.2 × 10^−16^). The centromere of five metacentric chromosomes were located after aligning six centromere-associated microsatellite markers to the assembly [71] (Table S4). Four of these five centromere regions co-localize with genomic windows enriched for repetitive elements (Figure 3d). Among repetitive elements, transposable elements (TEs) were the most common, and accounted for 36% of the assembled genome. Consistent with previous studies [46], the oyster genome is dominated by DNA transposons (32 % of the genome assembly) (Table 1), with *Helitrons* being the most abundant superfamily (Supplementary Figures S7-8).

**Figure 3.**
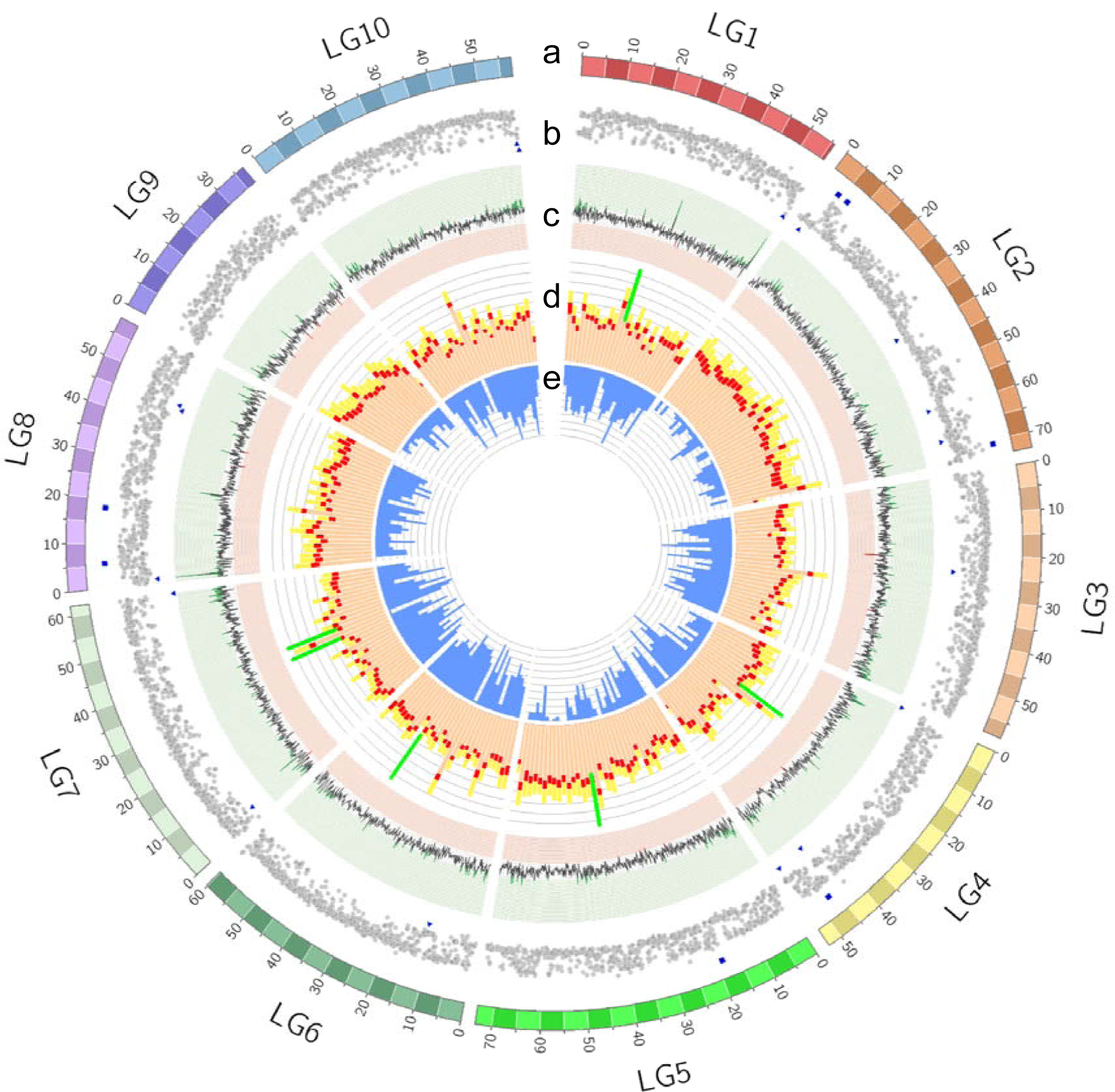
Circos plot depicting genome features across the 10 oyster chromosomes. (a) Oyster chromosomes (LG1–LG10 on a Mb scale). (b) short-read coverage plot. Coverage within 2SD of the mean are shown as grey circles. Abnormal sequence coverage (± 2SD form the mean) are indicated with a blue square or triangle, respectively. (c) GC content percentage (>35% in green; <31% in red). (d) Distribution of repeat elements: DNA transposons (light orange bar), retrotransposon TEs (red bar) and novel repeat elements (yellow bar). The location of centromeres is indicated with a green line. (e) Gene density (range 50-150). For tracks (b) and (c) a window size of 0.1 Mb was used, whereas for tracks (d) and (e) the size was increased to 0.2 Mb.

### Characterization of *Helitrons* in the Pacific oyster genome

*Helitrons* are rolling-circle transposable elements that have the ability to capture host gene fragments [72]. In maize, *Helitrons* have significantly influenced genome evolution, leading to genome variation among lines [73] and to a notable diversification of transcripts via exon shuffling of thousands of genes [74]. To refine the annotation of Pacific oyster *Helitrons*, a structure-based search [75] was performed in addition to the homology based approach described above. The localization of these elements was heterogeneous across the Pacific oyster chromosomes, with LG5 and LG8 showing a higher density of elements (>1SD above the average across chromosomes) (Figure S9). *Helitrons* in plant and animal genomes tend to accumulate in gene-poor regions [76]. However, this bias is less evident in *C. gigas*, with no significant association found between gene density and the number of *Helitrons* within a region. A comparison with other molluscan reference genome assemblies revealed that *C. gigas* had a remarkably high number of predicted *Helitron* related sequences (Figure 4).

**Figure 4.**
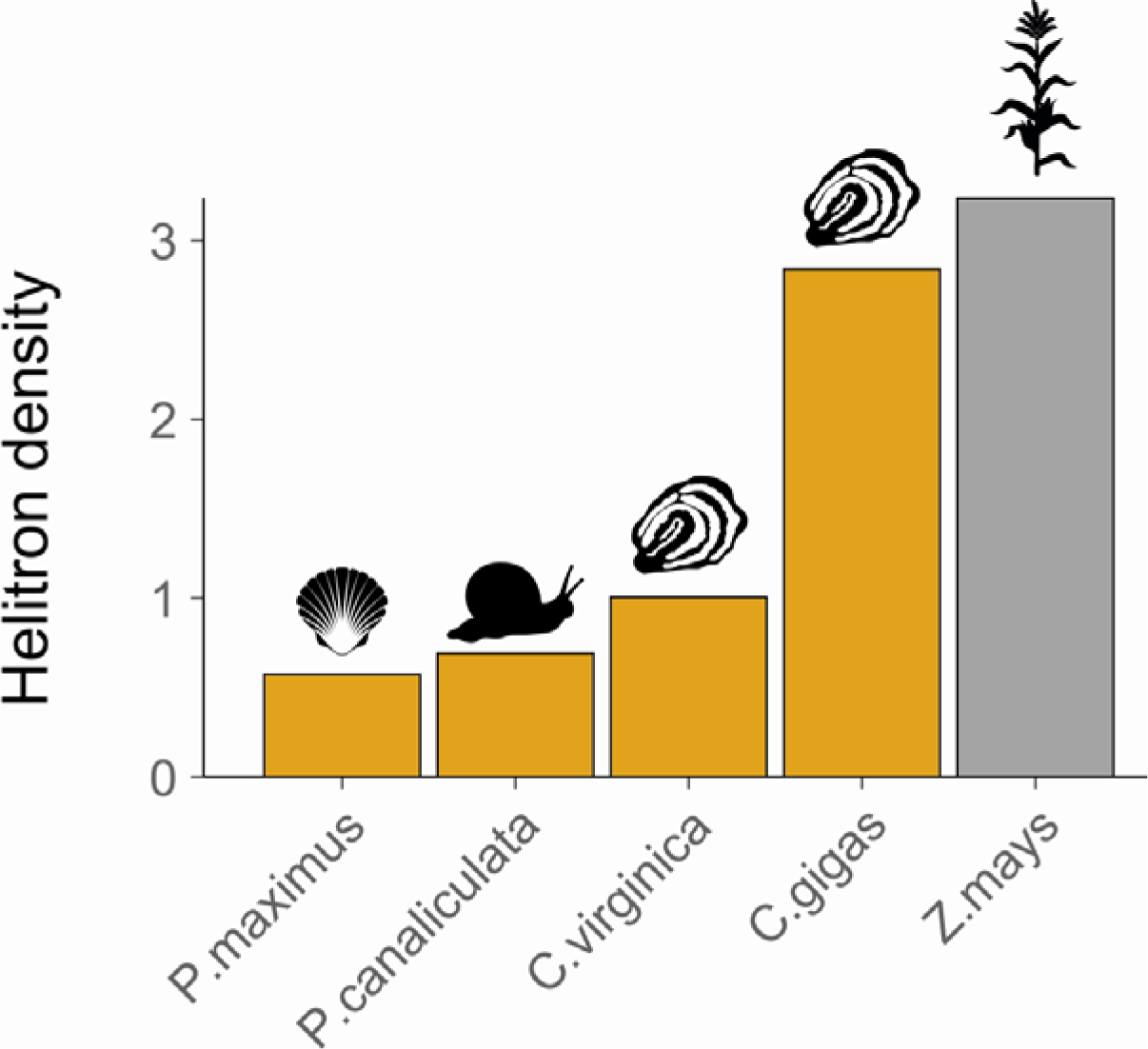
Density of *Helitrons* identified across four molluscan genomes (orange bars), including maize as a reference species (grey bar). The reference genome assembled for *C. gigas* was compared to the king scallop (*Pecten maximus*; GCF_902652985.1), golden apple snail (*Pomacea canaliculata*; GCF_003073045.1), and Atlantic oyster (*Crassostrea virginica*; GCF_002022765.2), with maize included as a reference species (*Zea mays*; GCF_000005005.2). *Helitron* density is expressed as the number of conserved 3’ ends over genome size (in Mb).

The Pacific oyster *Helitron-*like sequences possess the basic expected structure as observed in other taxa: TC sequence at the 5’ termini, CTAG motif on the 3’-terminus, and a 16-20 bp palindromic sequence that can form a hairpin structure upstream of the 3’-end. Likewise, they were also found to preferentially insert (86% of the cases) between the 5’-A and 3’-T nucleotides of the host AT target sites. Of the 751 intact *Helitrons* discovered through the *in silico* screening, 627 elements had a high 3’-end pairwise sequence similarity (identity of ∼85 % over 30 bp), suggesting they belong to the same family [76]. Notably, a significant fraction of these elements (261 out of 751) had sub-terminal inverted repeats, as revealed by a screening of their paired terminal ends with the Inverted Repeats Database (IRDB; https://tandem.bu.edu/cgi-bin/irdb/irdb.exe). This structural feature is characteristic of an alternative variant of *Helitrons* called *Helentrons*, which in its non-autonomous form known as HINE (Helentron-associated INterspersed Elements) has been recently associated to large numbers of satellite DNA in the oyster genome [77].

*Helitrons* have been observed to capture gene fragments in species such as maize and the little brown bat (*Myotis lucifugus*) [78, 79]. In *C. gigas*, a BLASTX [80] search against the UniRef database revealed that only 17 *Helitrons* (2%) carried gene fragments; alignment lengths >50 with at least 85% identity were considered a match. The Pacific oyster *Helitron*-like sequences were relatively short (mean = 1092 bp; SD = 557 bp), and lacked the main enzymatic hallmarks of autonomous elements (i.e., REP/Helicase domains). Non-autonomous *Helitrons* require the transposase expressed by their autonomous counterparts in order to amplify. Due to the fact this study did not detect evidence for the presence of autonomous mobile sequences in the Pacific oyster genome, these abundant *Helitron* elements are likely to be inactive, suggesting they are remnants of high levels of past activity in the evolutionary history of *C. gigas*.

## Conclusion

The new chromosome-level *C. gigas* genome assembly presented herein has a scaffold N50 of 58.4 Mb and a contig N50 of 1.8 Mb, representing a step advance on the previously published assembly, and will complement other high quality assemblies available or becoming available in the near-future. Approximately 30K putative protein coding genes were identified with an average of 3.9 transcripts per gene. DNA transposons dominated the repeat elements detected in the assembly, with *Helitrons* being found at a level substantially higher level than other molluscan species, suggesting their potential role in shaping the evolution of the *C. gigas* genome. The availability of a chromosome-level genome assembly is expected to support applied and fundamental research in this keystone ecological and aquaculture species.

## Availability of supporting data

Raw sequencing data has been submitted to the European Nucleotide Archive (ENA) under study accession number PRJEB35351. The genomic short read data are under accessions numbers ERX3728455, ERX3728453, ERX3728482, ERX3728546, ERX3728630 and ERX3728636; the raw reads of the Hi-C library are under accession numbers ERX3722775. PacBio Iso-Seq reads of pooled samples are available under accession numbers ERX3721883, ERX3722678 and ERX3722679. Raw PacBio reads from the nuclear DNA are available under accessions ERX3761471, ERX3761586, ERX3761587, ERX3761621, ERX3761714, ERX3761715, ERX3761720, ERX3762151, ERX3762342, ERX3762370, ERX3762371, ERX3762372 and ERX3762598. The Pacific oyster genome assembly is available at GenBank under accession number GCA_902806645.1.

## Abbreviations

bp: base pairs
BQ: base quality
BUSCO: Benchmarking Universal Single-Copy Orthologs
cM: centimorgan
cDNA: coding DNA
DNA: deoxyribonucleic acid
Gb: giga base pairs
GC: guanine-cytosine
Gb: gigabase pairs
kb: kilobase pairs
KEGG: Kyoto encyclopedia of genes and genomes
MAPQ: mapping quality
Mb: megabase pairs
N50: median size
PacBio: Pacific Biosciences
RNA: ribonucleic acid
RNA-Seq: RNA-sequencing
SMRT: single-molecule real-time

## Acknowledgements

The authors thank Katy Monteith, Darren Obbard and Carl Tucker for providing controls for the flow cytometry assay; Guernsey Sea Farms for rearing the female oyster used for generating the assembly; and Manu Kumar Gundappa and Richard Kuo from the Roslin Institute for their technical advice during the assembly and annotation steps. This work was supported by funding from the Natural Environment Research Council (NE/P010695/1) and Biotechnology and Biological Sciences Research Council (BB/S004343/1, BB/P013759/1 and BB/P013740/1). The cytogenetic mapping of BACs was supported by a grant from U.S. Department of Agriculture (2009-35205-05052).

## References

1. Salvi D, Macali A and Mariottini P. Molecular phylogenetics and systematics of the bivalve family Ostreidae based on rRNA sequence-structure models and multilocus species tree. PloS one. 2014;9 9:e108696–e. doi: 10.1371/journal.pone.0108696.

2. Salvi D and Mariottini P. Molecular taxonomy in 2D: a novel ITS2 rRNA sequence-structure approach guides the description of the oysters’ subfamily Saccostreinae and the genus Magallana (Bivalvia: Ostreidae). Zoological Journal of the Linnean Society. 2016;179 2:263–76. doi: 10.1111/zoj.12455.

3. FAO. 2020. Rome, Italy.

4. Wang H, Qian L, Liu X, Zhang G and Guo X. Classification of a Common Cupped Oyster from Southern China. Journal of Shellfish Research. 2010;29:857–66. doi: 10.2983/035.029.0420.

5. Robinson T, Griffiths C, Tonin A, Bloomer P and Hare M. Naturalized populations of oysters, Crassostrea gigas along the South African coast: Distribution, abundance and population structure. Journal of Shellfish Research. 2009;24:443–50. doi: 10.2983/0730-8000(2005)24[443:NPOOCG]2.0.CO;2.

6. Anglès d’Auriac MB, Rinde E, Norling P, Lapègue S, Staalstrøm A, Hjermann DØ, et al. Rapid expansion of the invasive oyster Crassostrea gigas at its northern distribution limit in Europe: Naturally dispersed or introduced? PLOS ONE. 2017;12 5:e0177481. doi: 10.1371/journal.pone.0177481.

7. Carrasco MF and Barón PJ. Analysis of the potential geographic range of the Pacific oyster Crassostrea gigas (Thunberg, 1793) based on surface seawater temperature satellite data and climate charts: the coast of South America as a study case. Biological Invasions. 2010;12 8:2597–607. doi: 10.1007/s10530-009-9668-0.

8. Miller PA, Elliott NG, Koutoulis A, Kube PD and Vaillancourt RE. Genetic Diversity of Cultured, Naturalized, and Native Pacific Oysters, Crassostrea Gigas, Determined from Multiplexed Microsatellite Markers. Journal of Shellfish Research. 2012;31 3:611–7, 7.

9. Meistertzheim A-L, Arnaud-Haond S, Boudry P and Thébault M-T. Genetic structure of wild European populations of the invasive Pacific oyster Crassostrea gigas due to aquaculture practices. Marine Biology. 2013;160 2:453–63. doi: 10.1007/s00227-012-2102-7.

10. Shatkin G, Shumway S and Hawes R. Considerations regarding the possible introduction of the Pacific oyster, Crassostrea gigas, to the Gulf of Maine: a review of global experience. Journal of Shellfish Research. 1997;16:463–78.

11. Jones MC, Dye SR, Pinnegar Jk, Warren R and Cheung WWL. Applying distribution model projections for an uncertain future: the case of the Pacific oyster in UK waters. Aquatic Conservation: Marine and Freshwater Ecosystems. 2013;23 5:710–22. doi: 10.1002/aqc.2364.

12. Wrange A-L, Valero J, Harkestad LS, Strand Ø, Lindegarth S, Christensen HT, et al. Massive settlements of the Pacific oyster, Crassostrea gigas, in Scandinavia. Biological Invasions. 2010;12 5:1145–52. doi: 10.1007/s10530-009-9535-z.

13. Herbert RJH, Humphreys J, Davies CJ, Roberts C, Fletcher S and Crowe TP. Ecological impacts of non-native Pacific oysters (Crassostrea gigas) and management measures for protected areas in Europe. Biodiversity and Conservation. 2016;25 14:2835–65. doi: 10.1007/s10531-016-1209-4.

14. Miossec L, Le Deuff R-M and Goulletquer P. Alien Species Alert: Crassostrea gigas (Pacific Oyster). ICES Cooperative Research Report. 2009;229:42.

15. Hedgecock D, Gaffney PM, Goulletquer P, Guo X, Reece K and Warr GW. The case for sequencing the Pacific oyster genome. Journal of Shellfish Research. 2005;24 2:429–41, 13.

16. Schwartz J, Réalis-Doyelle E, Dubos M-P, Lefranc B, Leprince J and Favrel P. Characterization of an evolutionarily conserved calcitonin signalling system in a lophotrochozoan, the Pacific oyster (Crassostrea gigas). The Journal of Experimental Biology. 2019;222 13:jeb201319. doi: 10.1242/jeb.201319.

17. Lafont M, Petton B, Vergnes A, Pauletto M, Segarra A, Gourbal B, et al. Long-lasting antiviral innate immune priming in the Lophotrochozoan Pacific oyster, Crassostrea gigas. Scientific Reports. 2017;7 1:13143. doi: 10.1038/s41598-017-13564-0.

18. Kocot KM. On 20 years of Lophotrochozoa. Organisms Diversity & Evolution. 2016;16 2:329–43. doi: 10.1007/s13127-015-0261-3.

19. Allen SK and Downing SL. Performance of triploid Pacific oysters, Crassostrea gigas (Thunberg). I. Survival, growth, glycogen content, and sexual maturation in yearlings. Journal of Experimental Marine Biology and Ecology. 1986;102 2:197–208. doi: https://doi.org/10.1016/0022-0981(86)90176-0.

20. Downing SL and Allen SK. Induced triploidy in the Pacific oyster, Crassostrea gigas: Optimal treatments with cytochalasin B depend on temperature. Aquaculture. 1987;61 1:1–15. doi: https://doi.org/10.1016/0044-8486(87)90332-2.

21. Guo X, DeBrosse GA and Allen SK. All-triploid Pacific oysters (Crassostrea gigas Thunberg) produced by mating tetraploids and diploids. Aquaculture. 1996;142 3:149–61. doi: https://doi.org/10.1016/0044-8486(95)01243-5.

22. Riviere G, Klopp C, Ibouniyamine N, Huvet A, Boudry P and Favrel P. GigaTON: an extensive publicly searchable database providing a new reference transcriptome in the pacific oyster Crassostrea gigas. BMC Bioinformatics. 2015;16 1:401. doi: 10.1186/s12859-015-0833-4.

23. Kim B-M, Kim K, Choi I-Y and Rhee J-S. Transcriptome response of the Pacific oyster, Crassostrea gigas susceptible to thermal stress: A comparison with the response of tolerant oyster. Molecular & Cellular Toxicology. 2017;13 1:105–13. doi: 10.1007/s13273-017-0011-z.

24. Yue C, Li Q and Yu H. Gonad Transcriptome Analysis of the Pacific Oyster Crassostrea gigas Identifies Potential Genes Regulating the Sex Determination and Differentiation Process. Marine biotechnology (New York, NY). 2018;20 2:206–19. doi: 10.1007/s10126-018-9798-4.

25. Feng D, Li Q, Yu H, Zhao X and Kong L. Comparative Transcriptome Analysis of the Pacific Oyster Crassostrea gigas Characterized by Shell Colors: Identification of Genetic Bases Potentially Involved in Pigmentation. PLOS ONE. 2015;10 12:e0145257. doi: 10.1371/journal.pone.0145257.

26. Zhang F, Hu B, Fu H, Jiao Z, Li Q and Liu S. Comparative Transcriptome Analysis Reveals Molecular Basis Underlying Fast Growth of the Selectively Bred Pacific Oyster, Crassostrea gigas. Frontiers in Genetics. 2019;10 610 doi: 10.3389/fgene.2019.00610.

27. Gutierrez AP, Bean TP, Hooper C, Stenton CA, Sanders MB, Paley RK, et al. A Genome-Wide Association Study for Host Resistance to Ostreid Herpesvirus in Pacific Oysters (Crassostrea gigas). G3: Genes|Genomes|Genetics. 2018;8 4:1273–80. doi: 10.1534/g3.118.200113.

28. Hedgecock D, Shin G, Gracey AY, Den Berg DV and Samanta MP. Second-Generation Linkage Maps for the Pacific Oyster Crassostrea gigas Reveal Errors in Assembly of Genome Scaffolds. G3 (Bethesda, Md). 2015;5 10:2007–19. doi: 10.1534/g3.115.019570.

29. Qi H, Song K, Li C, Wang W, Li B, Li L, et al. Construction and evaluation of a high-density SNP array for the Pacific oyster (Crassostrea gigas). PLOS ONE. 2017;12 3:e0174007. doi: 10.1371/journal.pone.0174007.

30. Gutierrez AP, Turner F, Gharbi K, Talbot R, Lowe NR, Peñaloza C, et al. Development of a Medium Density Combined-Species SNP Array for Pacific and European Oysters (Crassostrea gigas and Ostrea edulis). G3 (Bethesda). 2017;7 7:2209–18. doi: 10.1534/g3.117.041780.

31. Gutierrez AP, Matika O, Bean TP and Houston RD. Genomic Selection for Growth Traits in Pacific Oyster (Crassostrea gigas): Potential of Low-Density Marker Panels for Breeding Value Prediction. Frontiers in Genetics. 2018;9 391 doi: 10.3389/fgene.2018.00391.

32. Gutierrez AP, Symonds J, King N, Steiner K, Bean TP and Houston RD. Potential of genomic selection for improvement of resistance to ostreid herpesvirus in Pacific oyster (Crassostrea gigas). Animal Genetics. 2020;51 2:249–57. doi: 10.1111/age.12909.

33. Zhang G, Fang X, Guo X, Li L, Luo R, Xu F, et al. The oyster genome reveals stress adaptation and complexity of shell formation. Nature. 2012;490 7418:49–54. doi: 10.1038/nature11413.

34. Gomes-dos-Santos A, Lopes-Lima M, Castro LFC and Froufe E. Molluscan genomics: the road so far and the way forward. Hydrobiologia. 2020;847 7:1705–26. doi: 10.1007/s10750-019-04111-1.

35. Bolger AM, Lohse M and Usadel B. Trimmomatic: a flexible trimmer for Illumina sequence data. Bioinformatics. 2014;30 15:2114–20. doi: 10.1093/bioinformatics/btu170.

36. Marçais G and Kingsford C. A fast, lock-free approach for efficient parallel counting of occurrences of k-mers. Bioinformatics. 2011;27 6:764–70. doi: 10.1093/bioinformatics/btr011.

37. Dolezel J and Bartos J. Plant DNA Flow Cytometry and Estimation of Nuclear Genome Size. Annals of Botany. 2005;95 1:99–110. doi: 10.1093/aob/mci005.

38. Kajitani R, Toshimoto K, Noguchi H, Toyoda A, Ogura Y, Okuno M, et al. Efficient de novo assembly of highly heterozygous genomes from whole-genome shotgun short reads. Genome research. 2014;24 doi: 10.1101/gr.170720.113.

39. Ranallo-Benavidez TR, Jaron KS and Schatz MC. GenomeScope 2.0 and Smudgeplot for reference-free profiling of polyploid genomes. Nature Communications. 2020;11 1:1432. doi: 10.1038/s41467-020-14998-3.

40. Calcino AD, de Oliveira AL, Simakov O, Schwaha T, Zieger E, Wollesen T, et al. The quagga mussel genome and the evolution of freshwater tolerance. DNA Research. 2019;26 5:411–22. doi: 10.1093/dnares/dsz019.

41. Koren S, Walenz BP, Berlin K, Miller JR, Bergman NH and Phillippy AM. Canu: scalable and accurate long-read assembly via adaptive k-mer weighting and repeat separation. Genome Research. 2017; doi: 10.1101/gr.215087.116.

42. Chin C-S, Alexander DH, Marks P, Klammer AA, Drake J, Heiner C, et al. Nonhybrid, finished microbial genome assemblies from long-read SMRT sequencing data. Nature Methods. 2013;10 6:563–9. doi: 10.1038/nmeth.2474.

43. Walker BJ, Abeel T, Shea T, Priest M, Abouelliel A, Sakthikumar S, et al. Pilon: an integrated tool for comprehensive microbial variant detection and genome assembly improvement. PloS one. 2014;9 11:e112963–e. doi: 10.1371/journal.pone.0112963.

44. Li H. Minimap2: pairwise alignment for nucleotide sequences. Bioinformatics. 2018;34 18:3094–100. doi: 10.1093/bioinformatics/bty191.

45. Takeuchi T, Kawashima T, Koyanagi R, Gyoja F, Tanaka M, Ikuta T, et al. Draft Genome of the Pearl Oyster Pinctada fucata: A Platform for Understanding Bivalve Biology. DNA Research. 2012;19 2:117–30. doi: 10.1093/dnares/dss005.

46. Wang X, Xu W, Wei L, Zhu C, He C, Song H, et al. Nanopore Sequencing and De Novo Assembly of a Black-Shelled Pacific Oyster (Crassostrea gigas) Genome. Frontiers in Genetics. 2019;10 1211 doi: 10.3389/fgene.2019.01211.

47. Simao FA, Waterhouse RM, Ioannidis P, Kriventseva EV and Zdobnov EM. BUSCO: assessing genome assembly and annotation completeness with single-copy orthologs. Bioinformatics. 2015;31 19:3210–2. doi: 10.1093/bioinformatics/btv351.

48. Roach MJ, Schmidt SA and Borneman AR. Purge Haplotigs: allelic contig reassignment for third-gen diploid genome assemblies. BMC Bioinformatics. 2018;19 1:460. doi: 10.1186/s12859-018-2485-7.

49. Lieberman-Aiden E, van Berkum NL, Williams L, Imakaev M, Ragoczy T, Telling A, et al. Comprehensive mapping of long-range interactions reveals folding principles of the human genome. Science (New York, NY). 2009;326 5950:289–93. doi: 10.1126/science.1181369.

50. Putnam NH, O’Connell BL, Stites JC, Rice BJ, Blanchette M, Calef R, et al. Chromosome-scale shotgun assembly using an in vitro method for long-range linkage. Genome research. 2016;26 3:342–50. doi: 10.1101/gr.193474.115.

51. Thiriot-Quievreux C. Review of the literature on bivalve cytogenetics in the last ten years. Cahiers de Biologie Marine. 2002;43:17–26.

52. Li H and Durbin R. Fast and accurate short read alignment with Burrows-Wheeler transform. Bioinformatics (Oxford, England). 2009;25 14:1754–60. doi: 10.1093/bioinformatics/btp324.

53. English AC, Richards S, Han Y, Wang M, Vee V, Qu J, et al. Mind the Gap: Upgrading Genomes with Pacific Biosciences RS Long-Read Sequencing Technology. PLOS ONE. 2012;7 11:e47768. doi: 10.1371/journal.pone.0047768.

54. Warr A, Robert C, Hume D, Archibald AL, Deeb N and Watson M. Identification of Low-Confidence Regions in the Pig Reference Genome (Sscrofa10.2). Frontiers in Genetics. 2015;6 338 doi: 10.3389/fgene.2015.00338.

55. Durand NC, Robinson JT, Shamim MS, Machol I, Mesirov JP, Lander ES, et al. Juicebox Provides a Visualization System for Hi-C Contact Maps with Unlimited Zoom. Cell Syst. 2016;3 1:99–101. doi: 10.1016/j.cels.2015.07.012.

56. Krzywinski M, Schein J, Birol I, Connors J, Gascoyne R, Horsman D, et al. Circos: an information aesthetic for comparative genomics. Genome research. 2009;19 9:1639–45. doi: 10.1101/gr.092759.109.

57. Laetsch D and Blaxter M. BlobTools: Interrogation of genome assemblies. F1000Res, 2017.

58. Sedlazeck FJ, Rescheneder P, Smolka M, Fang H, Nattestad M, von Haeseler A, et al. Accurate detection of complex structural variations using single-molecule sequencing. Nature Methods. 2018;15 6:461–8. doi: 10.1038/s41592-018-0001-7.

59. Dobin A, Davis CA, Schlesinger F, Drenkow J, Zaleski C, Jha S, et al. STAR: ultrafast universal RNA-seq aligner. Bioinformatics. 2012;29 1:15–21. doi: 10.1093/bioinformatics/bts635.

60. Hoff KJ, Lomsadze A, Borodovsky M and Stanke M. Whole-Genome Annotation with BRAKER. Methods in molecular biology (Clifton, NJ). 2019;1962:65–95. doi: 10.1007/978-1-4939-9173-0_5.

61. Kuo RI, Cheng Y, Smith J, Archibald AL and Burt DW. Illuminating the dark side of the human transcriptome with TAMA Iso-Seq analysis. bioRxiv. 2019:780015. doi: 10.1101/780015.

62. Suzek BE, Wang Y, Huang H, McGarvey PB and Wu CH. UniRef clusters: a comprehensive and scalable alternative for improving sequence similarity searches. Bioinformatics. 2015;31 6:926–32. doi: 10.1093/bioinformatics/btu739.

63. Buchfink B, Xie C and Huson DH. Fast and sensitive protein alignment using DIAMOND. Nature Methods. 2015;12 1:59–60. doi: 10.1038/nmeth.3176.

64. Conesa A, Götz S, García-Gómez JM, Terol J, Talón M and Robles M. Blast2GO: a universal tool for annotation, visualization and analysis in functional genomics research. Bioinformatics. 2005;21 18:3674–6. doi: 10.1093/bioinformatics/bti610.

65. Xie C, Mao X, Huang J, Ding Y, Wu J, Dong S, et al. KOBAS 2.0: a web server for annotation and identification of enriched pathways and diseases. Nucleic Acids Research. 2011;39 suppl_2:W316–W22. doi: 10.1093/nar/gkr483.

66. https://www.ncbi.nlm.nih.gov/genome/annotation_euk/Crassostrea_gigas/102/ Accessed 01 September 2020.

67. RepeatMasker: http://www.repeatmasker.org. Accessed 17 April 2020.

68. Hubley R, Finn RD, Clements J, Eddy SR, Jones TA, Bao W, et al. The Dfam database of repetitive DNA families. Nucleic Acids Res. 2016;44 D1:D81–D9. doi: 10.1093/nar/gkv1272.

69. Bao W, Kojima KK and Kohany O. Repbase Update, a database of repetitive elements in eukaryotic genomes. Mobile DNA. 2015;6 1:11. doi: 10.1186/s13100-015-0041-9.

70. RepeatModeler: http://www.repeatmasker.org. Accessed April 17 2020.

71. Hubert S, Cognard E and Hedgecock D. Centromere mapping in triploid families of the Pacific oyster Crassostrea gigas (Thunberg). Aquaculture. 2009;288 3:172–83. doi: https://doi.org/10.1016/j.aquaculture.2008.12.006.

72. Kapitonov VV and Jurka J. Rolling-circle transposons in eukaryotes. Proceedings of the National Academy of Sciences. 2001;98 15:8714–9. doi: 10.1073/pnas.151269298.

73. Morgante M, Brunner S, Pea G, Fengler K, Zuccolo A and Rafalski A. Gene duplication and exon shuffling by helitron-like transposons generate intraspecies diversity in maize. Nature Genetics. 2005;37 9:997–1002. doi: 10.1038/ng1615.

74. Barbaglia AM, Klusman KM, Higgins J, Shaw JR, Hannah LC and Lal SK. Gene capture by Helitron transposons reshuffles the transcriptome of maize. Genetics. 2012;190 3:965–75. doi: 10.1534/genetics.111.136176.

75. Hu K, Xu K, Wen J, Yi B, Shen J, Ma C, et al. Helitron distribution in Brassicaceae and whole Genome Helitron density as a character for distinguishing plant species. BMC bioinformatics. 2019;20 1:354-. doi: 10.1186/s12859-019-2945-8.

76. Yang L and Bennetzen J. Structure-based discovery and description of plant and animal Helitrons. Proceedings of the National Academy of Sciences of the United States of America. 2009;106:12832–7. doi: 10.1073/pnas.0905563106.

77. Vojvoda Zeljko T, Pavlek M, Meštrović N and Plohl M. Satellite DNA-like repeats are dispersed throughout the genome of the Pacific oyster Crassostrea gigas carried by Helentron non-autonomous mobile elements. Scientific Reports. 2020;10 1:15107. doi: 10.1038/s41598-020-71886-y.

78. Yang L and Bennetzen JL. Distribution, diversity, evolution, and survival of Helitrons in the maize genome. Proc Natl Acad Sci U S A. 2009;106 47:19922–7. doi: 10.1073/pnas.0908008106.

79. Pritham EJ and Feschotte C. Massive amplification of rolling-circle transposons in the lineage of the bat Myotis lucifugus. Proceedings of the National Academy of Sciences. 2007;104 6:1895–900. doi: 10.1073/pnas.0609601104.

80. Altschul SF, Madden TL, Schäffer AA, Zhang J, Zhang Z, Miller W, et al. Gapped BLAST and PSI-BLAST: a new generation of protein database search programs. Nucleic Acids Res. 1997;25 17:3389–402. doi: 10.1093/nar/25.17.3389.

