## Supplementary Material for "A chromosome-level genome assembly for the Pacific oyster (*Crassostrea gigas*)"

**Supplemental Note A: Integration of the Pacific oyster genome assembly sequence with a cytogenetic map**

For fluorescence *in situ* hybridization (FISH), bacterial artificial chromosome (BAC) clones were first screened by PCR for microsatellites selected from a genetic linkage map (Hubert and Hedgecock 2004). BACs positive for mapped microsatellites from each linkage group were selected for FISH on metaphase chromosomes. BAC DNA was isolated with the Plasmid Midi kit (Qiagen), labelled with digoxigenin-11-dUTP by nick translation (Roche) and purified with the Wizard® DNA Clean-Up System (Promega) following the manufacturers’ instructions. Metaphase material was obtained from 6-hour old embryos and FISH was conducted following protocols described by Guo et al. (2007). Digoxigenin-labelled probes were detected with anti-digoxigenin-rhodamine (Roche). Chromosomes were then counterstained with 1.5 µg/ml 6-diamidino-2-phenylindole (DAPI) in antifade solution (Vector Laboratories). Oyster C0t-1 DNA was prepared according to the procedure described by Wang et al. (2005), and used to suppress repetitive sequences. Slides were observed using a Nikon E600 epifluorescence microscope equipped with a 3CCD camera (Optronics Laboratories). FISH signals were collected with appropriate filter set under 100x objective with the Image-Pro Plus software. Chromosomes were arranged and numbered from the longest to the shortest (Figure S4).

The complete sequences of the microsatellite clones were used for a NCBI BLASTN search (Zhang et al. 2000) against the chromosome level genome assembly (GenBank assembly accession GCA_902806645.1) using default parameters. The coordinates of the clones in the sequence assembly and their correspondence with the physical map are given in Table S1.

**Literature Cited**

Guo, X., Y. Wang, and Z. Xu, 2007 Genomic Analyses Using Fluorescence *In Situ* Hybridization, pp. 289-311 in *Aquaculture Genome Technologies*, edited by Z.J. Liu. Blackwell Publishing.

Hubert, S., and D. Hedgecock, 2004 Linkage maps of microsatellite DNA markers for the Pacific oyster *Crassostrea gigas*. *Genetics* 168 (1):351-362.

Wang, Y., Z. Xu, J.C. Pierce, and X. Guo, 2005 Characterization of Eastern Oyster (*Crassostrea virginica* Gmelin) Chromosomes by Fluorescence *In Situ* Hybridization with Bacteriophage P1 Clones. *Marine Biotechnology* 7 (3):207-214.

**Supplemental Note B: Oyster genome annotation**

The oyster genome annotation was based on two sources of data: PacBio long-read IsoSeq data generated for this study and thirteen, previously released, paired-end RNA-seq datasets Zhang (Zhang et al. 2012). Short-read data was mapped with STAR v.2.5.1b (Dobin et al. 2012) with parameters ‘--outFilterType BySJout --outSAMunmapped Within --outSAMtype BAM SortedByCoordinate --outSAMattrIHstart 0 --outFilterIntronMotifs RemoveNoncanonical --alignSoftClipAtReferenceEnds No’. Transcript models were created by BRAKER v.2.1.5 (Hoff et al. 2019) with GeneMark v.4.61_lic (Lomsadze et al. 2005) and Augustus v3.3.3 (Stanke et al. 2008) using only the paired-end RNA-seq datasets (Table S3). The Iso-Seq CCS reads were mapped with minimap2 v.2.16 (Li 2018) with options ‘-ax splice -uf --junc-bed’ using splice-junction information from the mapped RNA-seq reads, keeping only those junctions which had a support of at least 5 uniquely mapped reads.

The Iso-Seq transcript models were processed by TAMA package (Kuo et al. 2019). The mapped sequences were first collapsed with tama_collapse: tama_collapse.py -s input.sam -f cgigas_uk_roslin_v1.softmasked.simple.fa -p input.collapse -x capped -sjt 20 -lde 2 -a 100 -z 100 and the sample specific models were then merged together with the tama_merge.py. Read support of the IsoSeq models were calculated by tama_read_support_level.py and the transcript models containing poly-A sequences were obtained with tama_remove_polya_models_levels.py.

Finally, the IsoSeq and RNA-seq based transcript models were merged together by tama_mereg.py. Protein-coding transcripts and translation start and end positions were predicted by mapping known protein sequences from UniRef90 (Suzek et al. 2015) to the oyster transcripts by Diamond v.0.9.31 (Buchfink et al. 2015) using default parameters. If the predicted start and end codons were non canonical then the transcript was parsed 50 codon upstream and downstream, observing the reading frame, for canonical start and stop codons and if found the coding sequence was extended. Models that contained a frameshift within the coding sequence were classified as pseudogenes.

**Literature cited**

Buchfink, B., C. Xie, and D.H. Huson, 2015 Fast and sensitive protein alignment using DIAMOND. *Nature Methods* 12 (1):59-60.

Dobin, A., C.A. Davis, F. Schlesinger, J. Drenkow, C. Zaleski *et al.*, 2012 STAR: ultrafast universal RNA-seq aligner. *Bioinformatics* 29 (1):15-21.

Hoff, K.J., A. Lomsadze, M. Borodovsky, and M. Stanke, 2019 Whole-Genome Annotation with BRAKER. *Methods Mol Biol* 1962:65-95.

Kuo, R.I., Y. Cheng, J. Smith, A.L. Archibald, and D.W. Burt, 2019 Illuminating the dark side of the human transcriptome with TAMA Iso-Seq analysis. *bioRxiv*:780015.

Li, H., 2018 Minimap2: pairwise alignment for nucleotide sequences. *Bioinformatics* 34 (18):3094-3100.

Lomsadze, A., V. Ter-Hovhannisyan, Y.O. Chernoff, and M. Borodovsky, 2005 Gene identification in novel eukaryotic genomes by self-training algorithm. *Nucleic acids research* 33 (20):6494-6506.

Stanke, M., M. Diekhans, R. Baertsch, and D. Haussler, 2008 Using native and syntenically mapped cDNA alignments to improve de novo gene finding. *Bioinformatics* 24 (5):637-644.

Suzek, B.E., Y. Wang, H. Huang, P.B. McGarvey, and C.H. Wu, 2015 UniRef clusters: a comprehensive and scalable alternative for improving sequence similarity searches. *Bioinformatics* 31 (6):926-932.

Zhang, G., X. Fang, X. Guo, L. Li, R. Luo *et al.*, 2012 The oyster genome reveals stress adaptation and complexity of shell formation. *Nature* 490 (7418):49-54.

Zhang, Z., S. Schwartz, L. Wagner, and W. Miller, 2000 A greedy algorithm for aligning DNA sequences. *J Comput Biol* 7 (1-2):203-214.

**Table S1. Correspondence between the ten *C. gigas* linkage groups (LG) and chromosomes (Chr)**

| Assembly LG | Chr | BAC ID | Seq-Tag accession N˚ | FISH location | cgigas_uk_roslin_v1  coordinates (bp) |
| --- | --- | --- | --- | --- | --- |
| 1 | 7 | 091-E16 | AF201464.1 | 7p | 8724555 to 8724997 |
| 2 | 1 | 049-N13 | AF051172.1 | 1q | 30621939 to 30622151 |
| 3 | 9 | 062-L19 | AF468559.1 | 9q | 24424135 to 24424649 |
| 4 | 6 | 001-I13 | AF468596.1 | 6p | 5612565 to 5613172; 5906743 to 5907363 |
| 5 | 3 | 001-L16 | AF468530.1 | 3q | 1598348 to 1598748 |
| 6 | 2 | 108-I23 | AB091583.1 | 2q | 46504621 to 46505330 |
| 7 | 4 | 089-J17 | AF468536.1 | 4q | 18867646 to 18868016 |
| 8 | 5 | 171-N24 | AY999705.1 | 5q | 25208347 to 25208807 |
| 9 | 10 | 184-F06 | AF468582.1 | 10q | 29057540 to 29058059; 29137829 to 29138312 |
| 10 | 8 | 004-P05 | DQ002500.1 | 8q | 36564944 to 36565348 |

**Table S2. Validation of the *C. gigas* genome based on the alignment of Pac-Bio long reads**

|  | **Structural assembly errors** | | | |
| --- | --- | --- | --- | --- |
| **Assembly** | **DEL** | **DUP** | **INV** | **INS** |
| cgigas_uk_roslin_v1 | 7,441 | 440 | 214 | 2,779 |

**Table S3 Paired-end RNA-seq read information.** Man1= outer edge of mantle; Dg = digestive gland; Fgo = female gonad; Gill = gill; Man2 = inner part of mantle; Amu = adductor muscle; Hem = hemocyte; Lpa = labial palp; Mgo = male gonad; G3 = adult (undefined tissue); early = early developmental stages; late = late planktonic stages and spat; Rem = mixture of adult tissue (hemolymph, mantle, gonad, digestive gland, gill, labial palp, and adductor muscle).

| Sequence Read Archive ID | Experiment ID | Experiment run ID | Tissue | Uniquely mapped reads (%) | Total mapped reads (%) |
| --- | --- | --- | --- | --- | --- |
| SRA045614 | SRX093411 | SRR334212 | Man1 | 80.31 | 86.73 |
| SRA045614 | SRX093412 | SRR334213 | Dgl | 79.29 | 85.46 |
| SRA045614 | SRX093413 | SRR334214 | Fgo | 78.08 | 83.6 |
| SRA045614 | SRX093414 | SRR334215 | Gill | 78.61 | 85.75 |
| SRA045614 | SRX093415 | SRR334216 | Man2 | 81.63 | 87.84 |
| SRA045614 | SRX093416 | SRR334217 | Amu | 83.62 | 88.11 |
| SRA045614 | SRX093417 | SRR334218 | Hem | 80.58 | 86.08 |
| SRA045614 | SRX093418 | SRR334219 | Lpa | 77.76 | 84.42 |
| SRA045614 | SRX093419 | SRR334220 | Mgo | 83.21 | 89.17 |
| SRA045614 | SRX093420 | SRR334221 | G3 | 80.35 | 84.96 |
| SRA045614 | SRX093538 | SRR334339 | early | 77.37 | 82.72 |
| SRA045614 | SRX093539 | SRR334340 | late | 73.36 | 78.23 |
| SRA056342 | SRX170733 | SRR526975 | Rem | 78.44 | 84.84 |

**Table S4. Microsatellite markers associated with *C.gigas* centromeres**

| LG ID | Locus | GenBank accession N˚ |
| --- | --- | --- |
| LG6 | *uscCg205* | AY999703 |
| LG7 | *ucdCg028* | AF051178 |
| LG7 | *ucdCg197* | AF468595 |
| LG1 | *imbCg44* | Y12085 |
| LG4 | *imbCg049* | Y12086 |
| LG5 | *ucdCg137* | AF468541 |

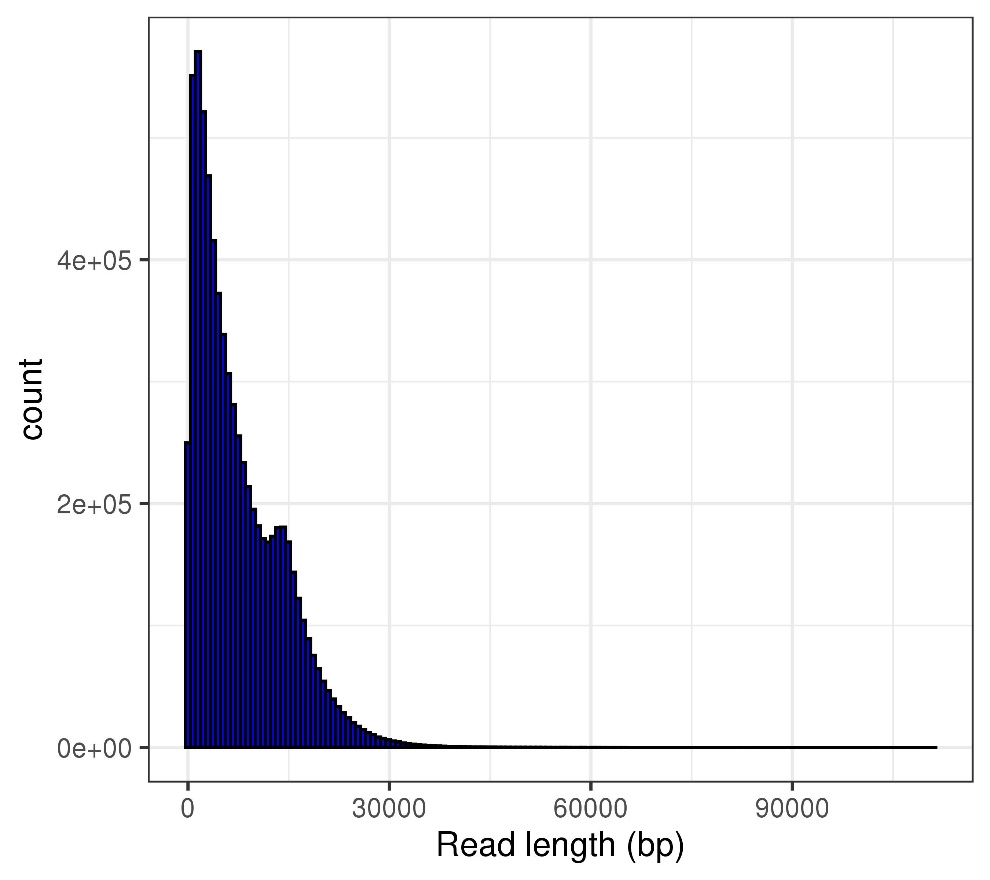

**Figure S1. Read length distribution of raw PacBio reads**

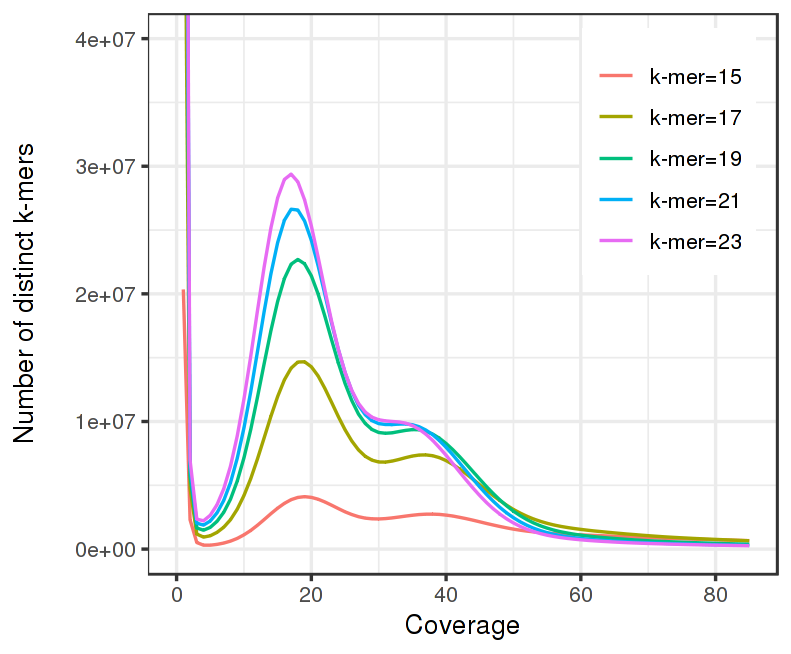

**Figure S2. Distribution of different k-mer depths estimated from short read data.** Two peaks at ~19 and ~37 are observed, suggesting the Pacific oyster genome is highly heterozygous.

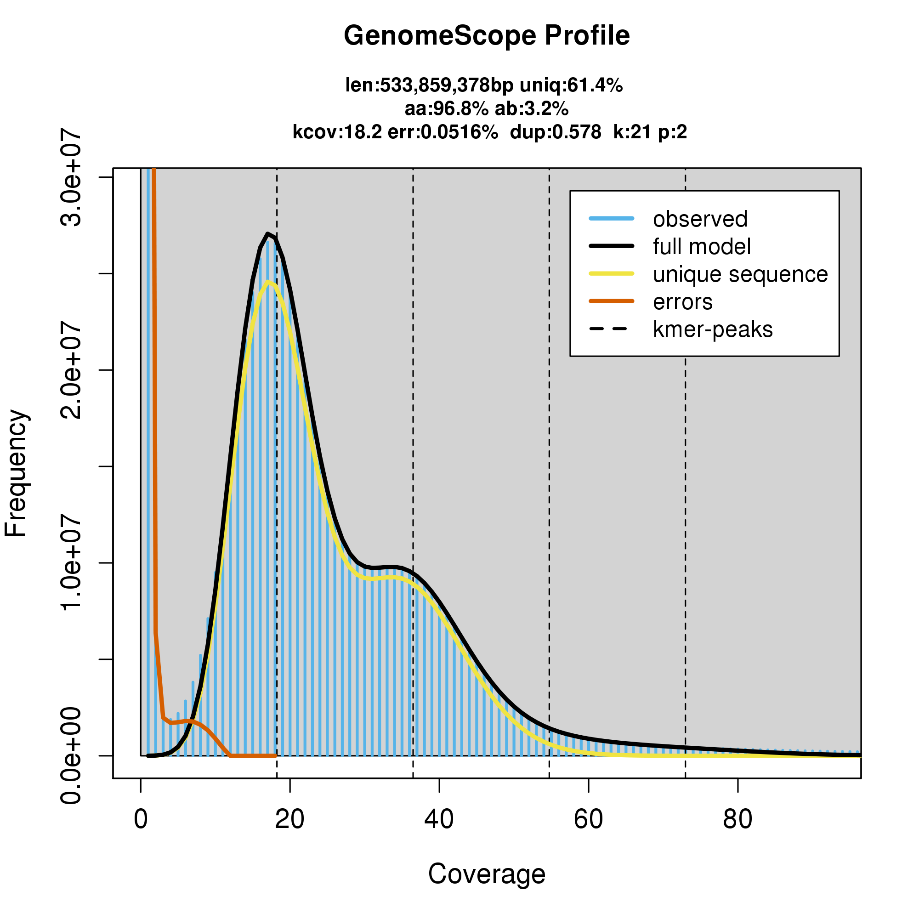

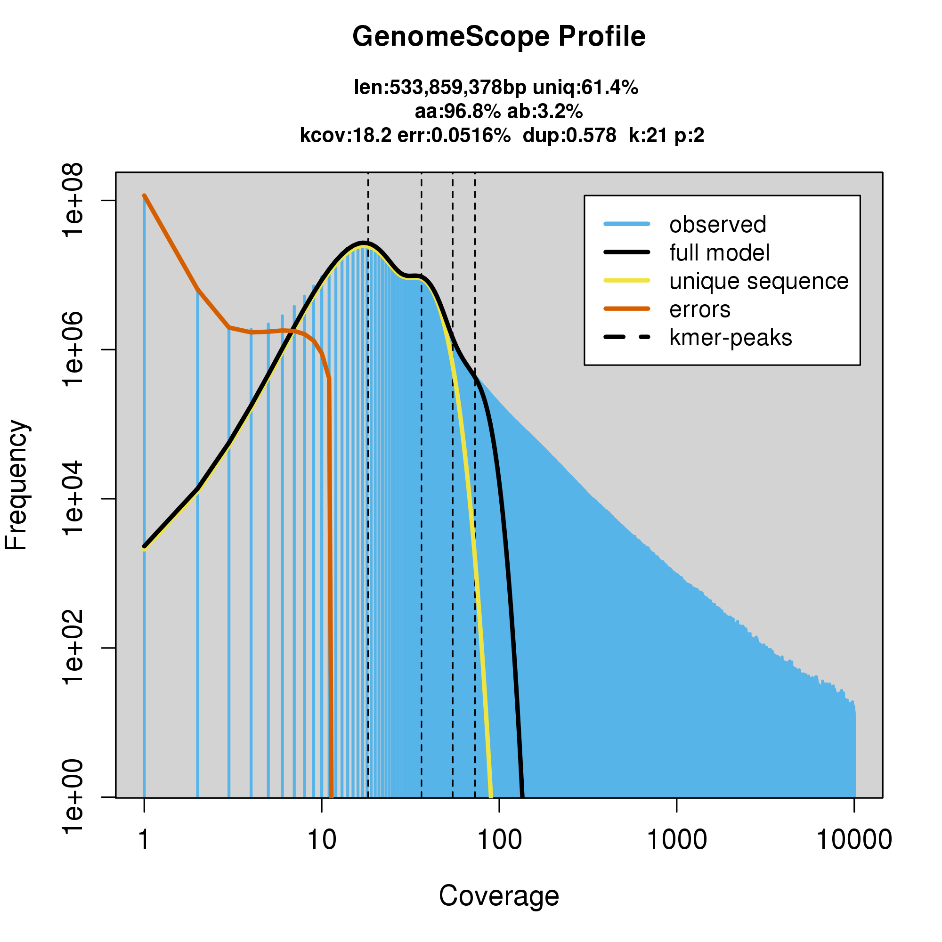

**Figure S3. GenomeScope results plots of the 21-mer k-mer content of *C. gigas***

**Figure S4. A karyotype of Pacific oyster with each chromosome identified by FISH with a sequence-tagged BAC clone.** Sequence tag accession numbers are given in Table S1.

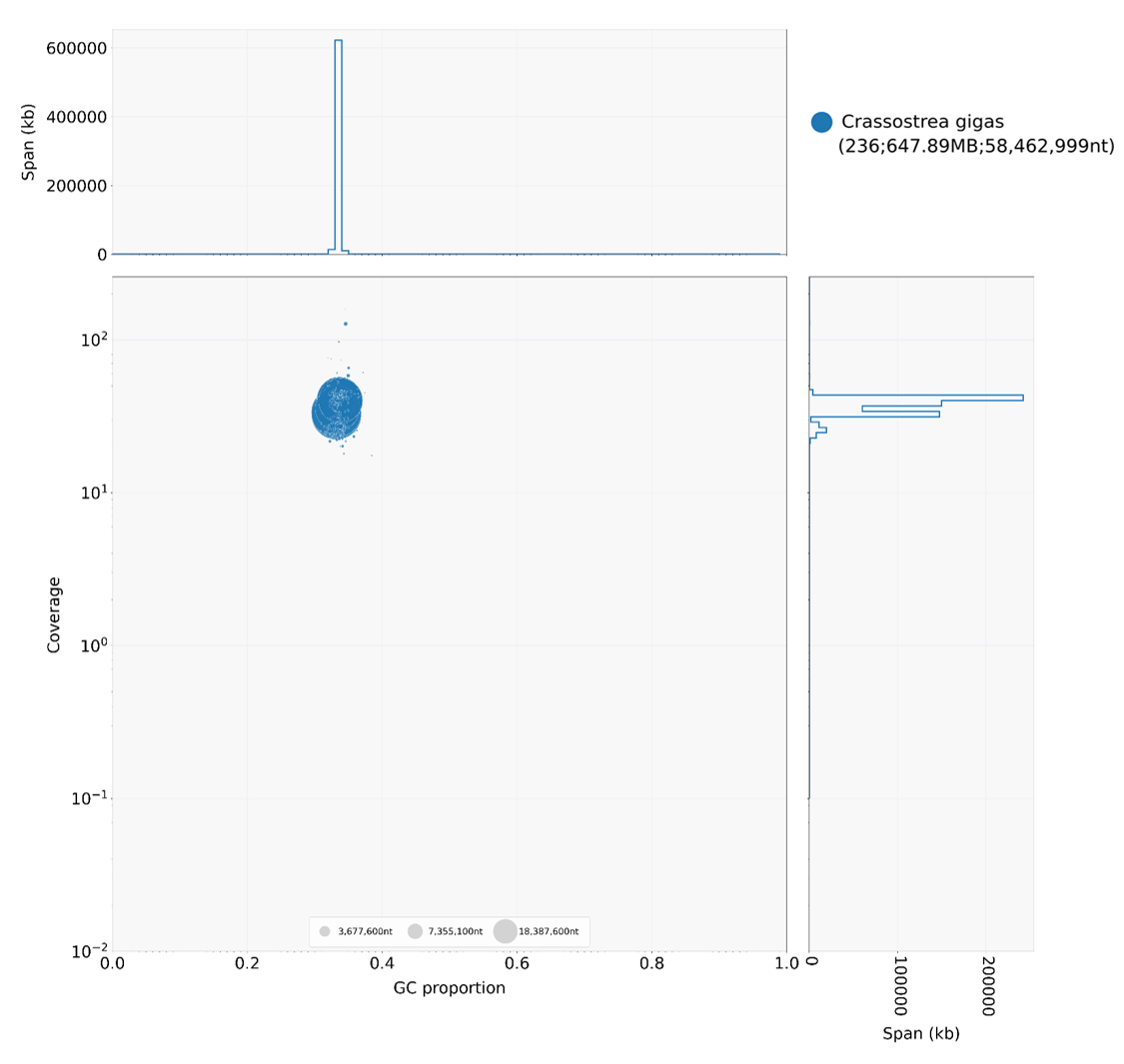

**Figure S5. Taxon-annotated GC-coverage plot for the Pacific oyster genome assembly.** Central panel: The GC-content (x-axes) of each scaffold/contig is plotted against their average short read coverage (y-axes). Each scaffold/contig in the assembly is represented by a circle, which is colour coded according to the best match to taxonomically annotated sequence databases (upper right legend). The circles are scaled to scaffold/contig length according to the legend on the bottom section of the panel. Right panel: Nucleotide span in kb at each coverage level. Top panel: Nucleotide span in kb at each GC proportion.

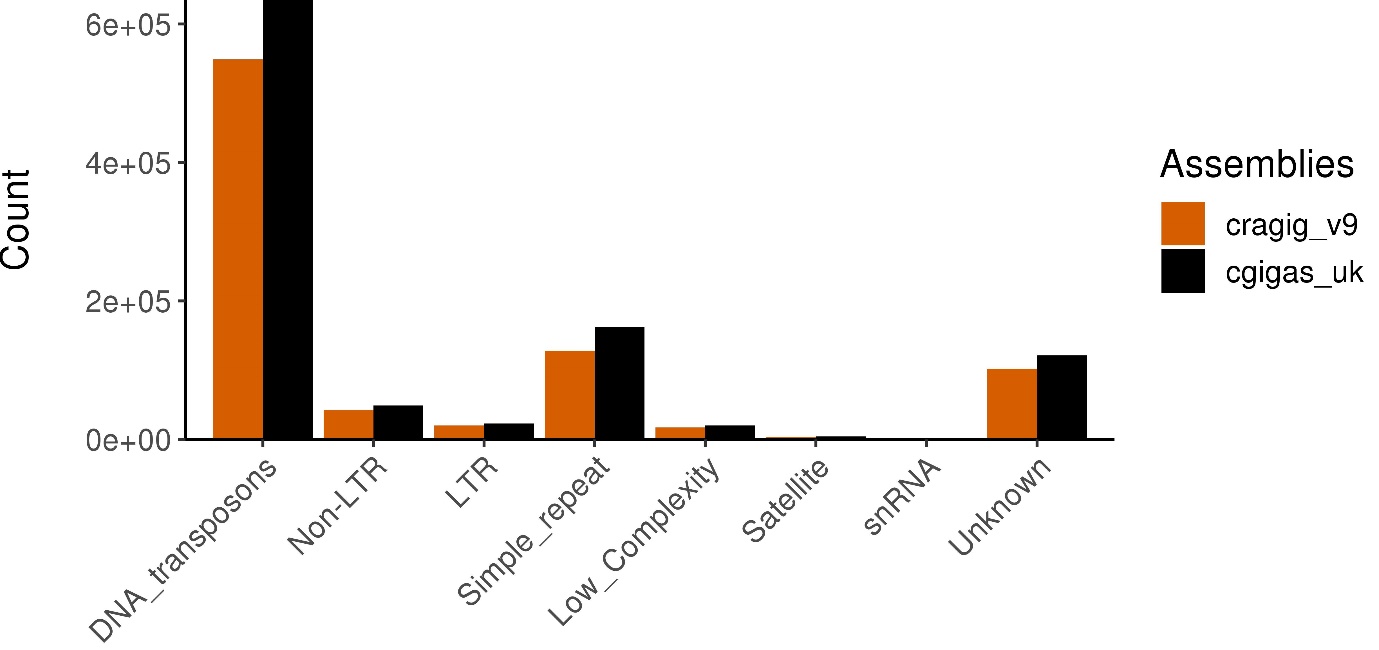

**Figure S6. Comparison of major categories of repeat elements observed within the previous genome assembly (in orange) and the improved reference genome assembly (in black).**

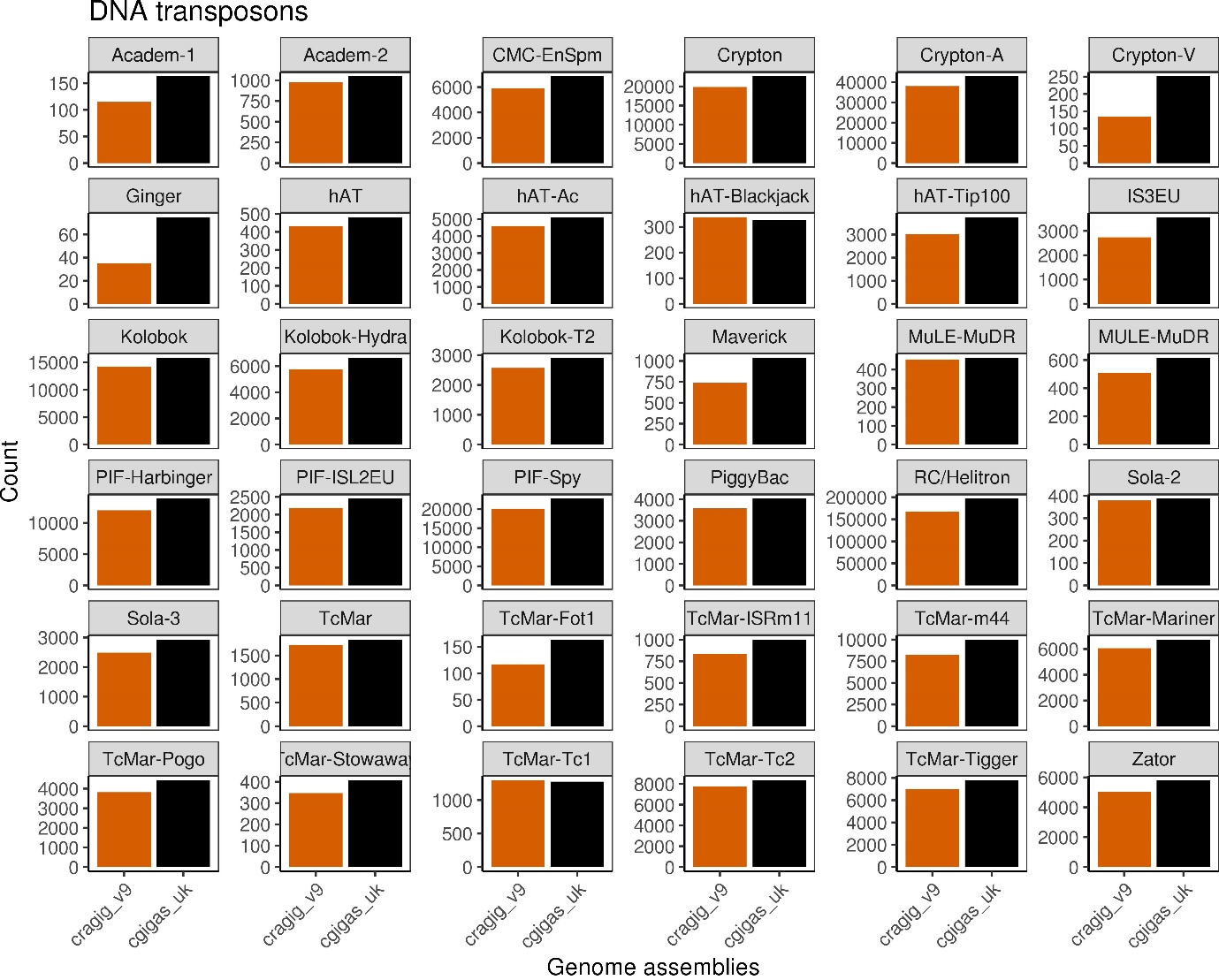

**Figure S7. Number of DNA transposons identified in the previous oyster genome assembly (in orange) and the improved assembly (in black), based on a RepeatModeler and RepeatMasker analysis**

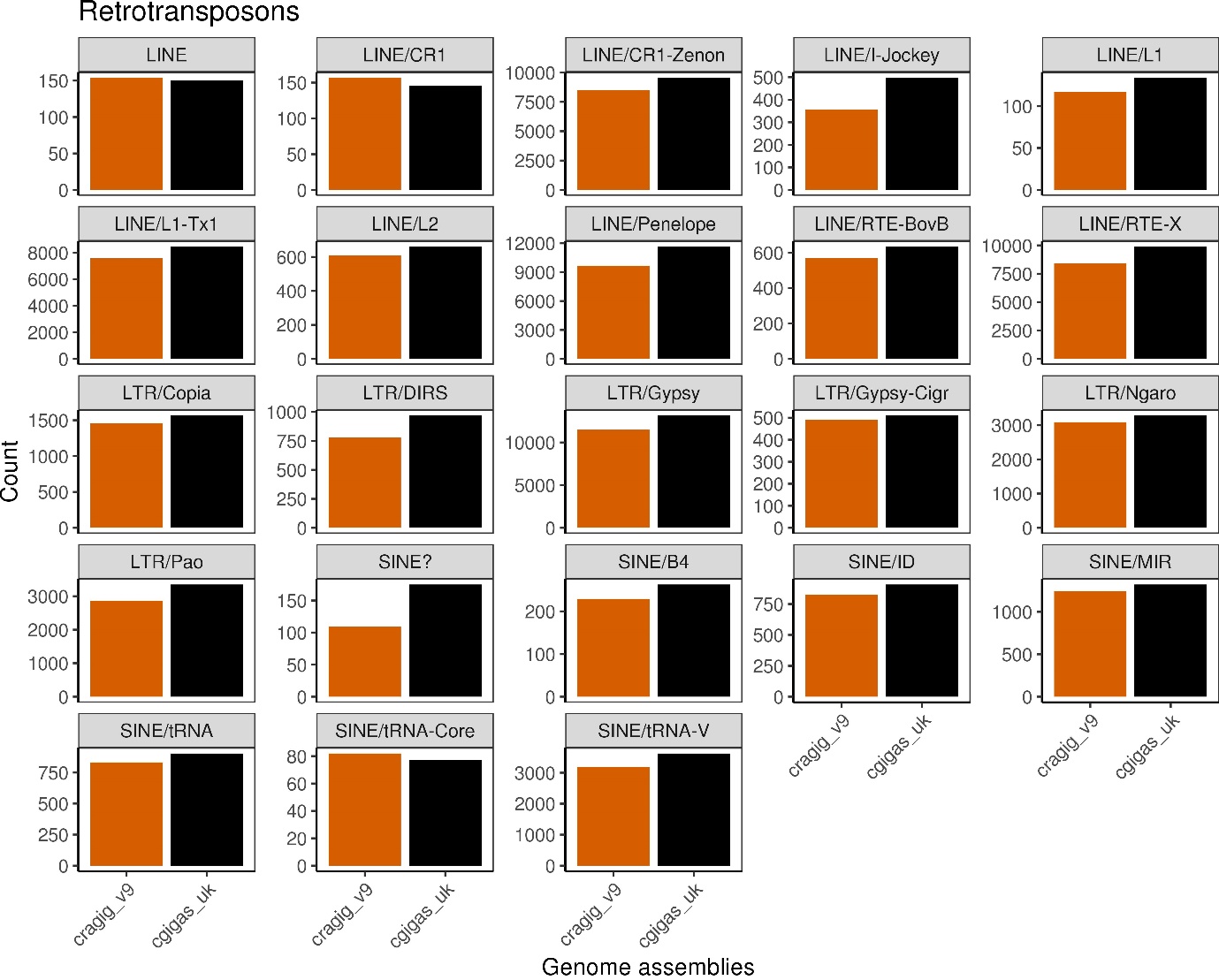

**Figure S8. Number of retrotransposon TEs identified in the previous oyster genome assembly (in orange) and the improved assembly (in black), based on a RepeatModeler and RepeatMasker analysis**

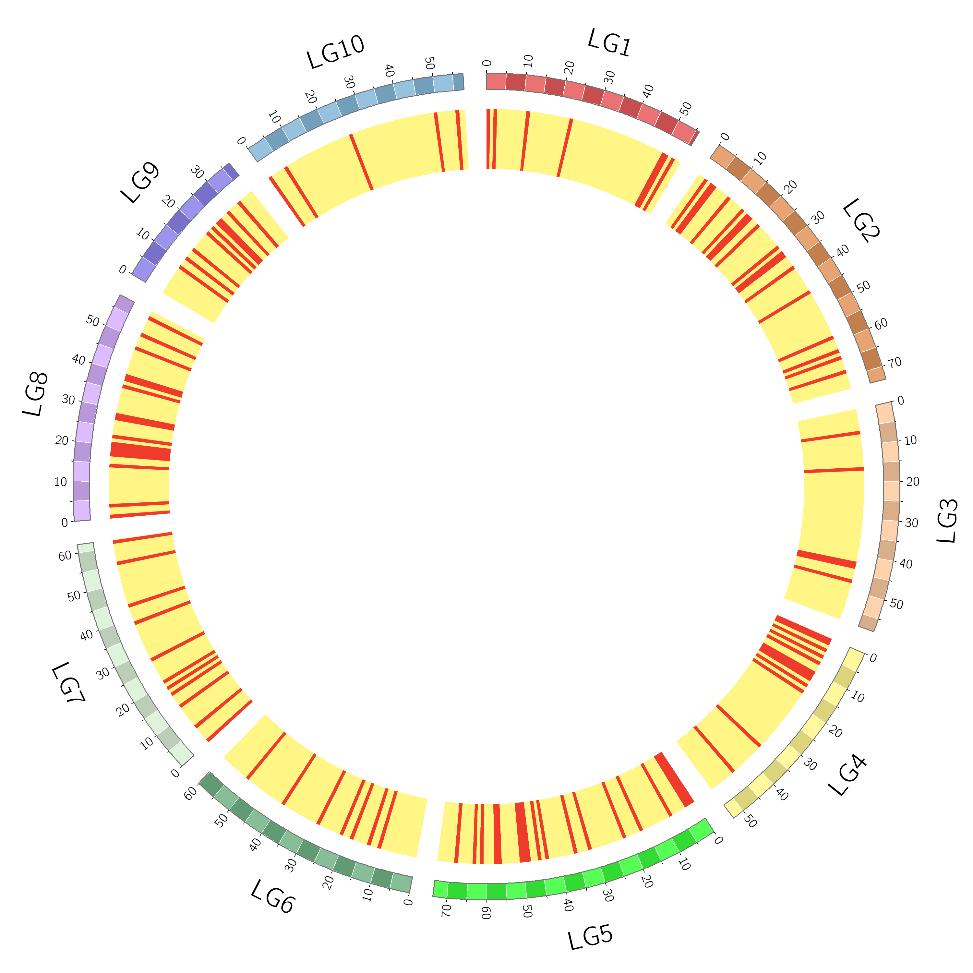

**Figure S9. Location of putative *Helitrons* along the ten chromosomes of the improved reference genome assembly for the Pacific oyster**
